# Revealing pH-dependence and independence of the characteristics of a *β* sheet-forming antimicrobial *peptide*^†^

**DOI:** 10.1101/2025.10.02.679978

**Authors:** M.N. Hamidabad, R.A. Mansbach

**Affiliations:** Physics Department, Concordia University, Montréal, QC, H4B 1R6, Canada

## Abstract

Membrane-active antimicrobial peptides (AMPs) are a promising potential solution to combat rising antimicrobial resistance (AMR) due to their selective interaction with negatively charged bacterial membranes, but their behavior is controlled by their charge states, which in turn depend on the local pH in which they find themselves. In this study, we employ constant pH molecular dynamics (CpHMD) simulations to investigate the pH-dependent behavior of a 13-residue-long positively charge AMP, GL13K, focusing on the deprotonation states of lysine residues in a single GL13K AMP and their impact on its structural dynamics. We determine *pK_a_* values of the critical lysine residues and show that the last lysine located near the C-terminus (LYS11) has a significant deprotonation difference with other lysine residues. We observe that increasing the pH results in changes in the metastable conformational states including collapse of the peptide and highlight the stabilization of a potentially therapeutically-relevant *β* hairpin configuration in pH levels leading to partial protonation of the lysines. Overall, our study shows the pH-dependent conformational dynamics and *pK_a_* variations of lysine residues in the GL13K antimicrobial peptide, providing critical insights into its structural behavior in solution. These findings establish a necessary rigorous foundation for further exploration of GL13K in more complex systems, advancing its potential development as an antimicrobial agent.

## 1 Introduction

Antimicrobial peptides (AMPs) are a diverse group of short proteins integral to the innate immune system that can also be designed and developed synthetically as drug candidates with desirable properties ^1^. They exhibit broad-spectrum activity against a variety of bacteria via several different mechanisms, including direct membrane disruption ^2^. The efficacy of AMPs is associated with their physicochemical properties, particularly their hydrophobicity and charge state, which can be significantly influenced by environmental factors such as local ion concentration and local pH. Additionally, the mechanisms of action of membrane-disruptive AMPs are directly linked to their available conformations in the vicinity of the pathogenic membrane, which is likewise highly dependent on their charge states ^3^ and thus on the local environment imposed by the membrane. The potential interrelation between the charge state of an AMP and its conformational structure is therefore a critical area for investigation. On the one hand, the charge state may significantly alter the conformational ensemble of the AMPs; on the other hand, the conformations control the intra-molecular interactions, which may change the local environment surrounding a residue, resulting in e.g. different *pK*_*a*_ values of residues of the same type ^4^.

In this work, we focus on characterizing pH-dependent charge and conformational states of the synthetic 13-residue-long AMP, GL13K (sequence *GKIIKLKASLKLL*, chemical structure Fig. 1). This AMP inhibits the gram-negative endotoxin, lipopolysaccharide (LPS), and exhibits both Gram-negative and Gram-positive bactericidal activity while maintaining low hemolyticity ^5^. It was derived from the salivary protein parotid secretory protein (BPIFA2, formerly known as PSP) by isolating residues 141 to 153 and replacing glutamine, asparagine, and aspartic acid in positions 2, 5, and 11 with lysine residues, which have a standard *pK*_*a*_ of 10.5 in aqueous solution. These substitutions confer upon the peptide a strong pH dependence and increase the net charge of the peptide from +1 to +5 at physiological pH (7.4). The pH dependence of the peptide’s charge state leads to pH-dependent conformational and aggregative behavior, which appears linked to its capacity to disrupt membranes of both Gram-negative and Gram-positive bacteria, while not impacting host red blood cell membranes. Specifically, at neutral pH, in solution, GL13K predominantly takes on random coil conformations, while at high pH, it shows an increasing propensity towards *β* sheet formation ^6,7^. Similarly, it has been observed that near eukaryotic membrane mimics, such as those modeled by DOPC liposomes, it adopts primarily random-coil conformations, while, in the presence of gram-negative inner membrane/gram-positive plasma membrane mimics, such as those modeled by negatively charged DOPG liposomes, it adopts *β* sheet structures ^8^.

**Fig. 1.**
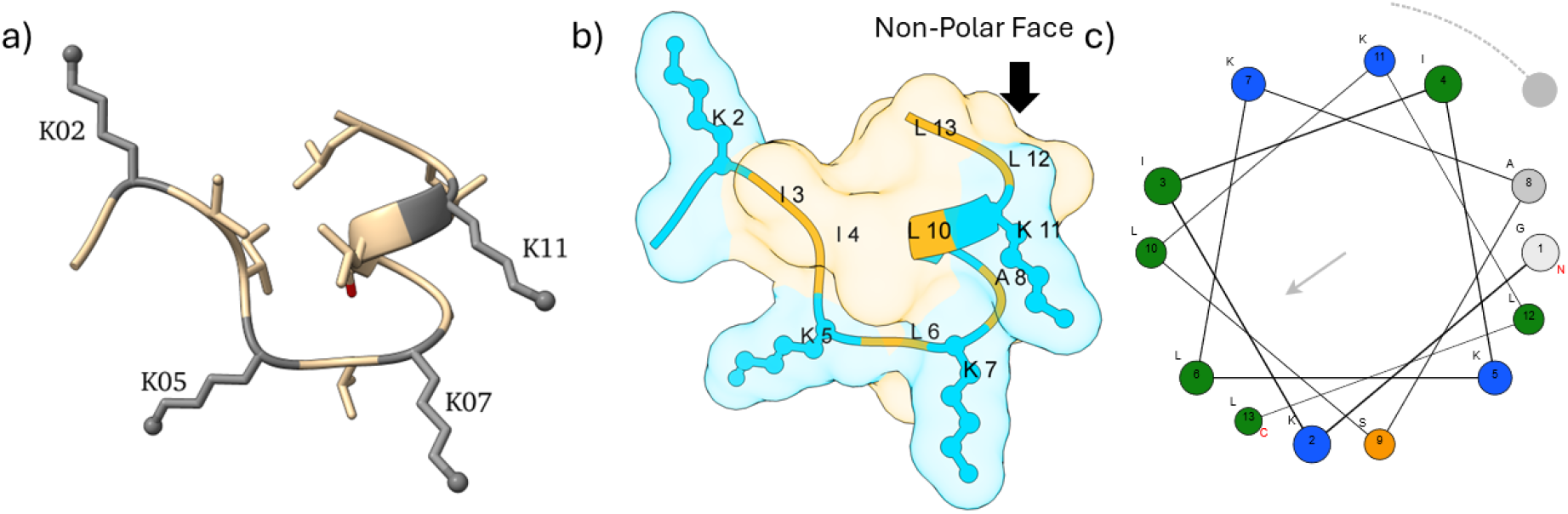
Structure of GL13K and location of titratable residues. (a) Initial structure of GL13K peptide predicted by PEP-FOLD4 web-server ^21^. Lysine residues are colored in gray with the side-chain Nitrogen (Nitrogen of *ε*-amino group as the titratable site) represented by a black ball. (b) The hydrophobic and hydrophilic surfaces of this structure are colored in gold and blue, respectively. The hydrophobic amino acids and Lysine residues are labeled. Balls and sticks represent lysine side chains. (c) Helical wheel of GL13K is plotted with charged polar residues in blue, uncharged polar residues in orange, hydrophobic residues with bulky side-chains in green, and small non-polar residues in light gray ^22^. Subplots (a) and (b) were rendered with UCSF ChimeraX ^23^, developed by the Resource for Biocomputing, Visualization, and Informatics at the University of California, San Francisco, with support from National Institutes of Health R01-GM129325 and the Office of Cyber Infrastructure and Computational Biology, National Institute of Allergy and Infectious Diseases.

Because the conformational changes GL13K undergoes are accompanied by aggregative behavior (non-interacting random coil to strongly aggregative *β* sheet formation), both conformational changes and self-assembly propensity appear linked to its disruptive mechanisms, which appear to be a combination of pore-forming and micellization ^6,8^. It has the ability to self-assemble into multiple types of supramolecular structures ^9^. More precisely, while GL13K can form some crystalline *β* -sheets at an interface even at pH 7.4, it does not self-assemble into fibrils under these conditions ^10^; it only forms fibrils at higher pH levels in solution ^11^ and when interacting with lipopolysaccharide, lipoteichoic acid (components of Gram-positive bacterial cell walls), or the cell wall of Gram-positive bacteria ^12^. It has been suggested that the lysine residue located on the non-polar face (Figure 1-b, LYS11) controls fibril formation through the inhibition of peptide self-association when charged (at neutral pH) and the removal of this inhibition at higher pH, when it becomes uncharged ^10^.

Taken all together, this means that a crucial piece of groundwork in a better understanding of the complex process of GL13K conformational shift and aggregative behavior is a comprehensive calculation of the pH-dependence of its lysine residues and how they control its single-peptide conformations. Such single-molecule details are remarkably difficult to capture experimentally, which leads to a computational approach. Molecular dynamics (MD) simulations are a powerful tool for the study of peptides, such as molecular modeling of peptides ^13^, peptide-eukaryotic membrane interaction studies ^14^, peptide-prokaryotic membrane interaction studies ^15^, and designing new antimicrobial and anticancer peptides ^16^. Although MD simulation is a powerful computational tool, it has some considerable limitations; in traditional MD simulations, the charge state of all amino acids in the sequence is set to the most probable value at the beginning of the simulation and during the simulations is fixed ^17^. This limitation motivated us to use constant pH molecular dynamics to investigate the effects of pH on GL13K, in particular the role of lysine residues as titratable sites at different pH levels.

Constant pH molecular dynamics (CpHMD) is an advanced computational method used in molecular dynamics simulations to model pH-dependent processes in biological systems ^18^. The most common method of implementing it utilizes the *λ* -dynamics technique to represent the dynamic protonation states of a titratable site using a continuous variable (*λ* ), enabling sampling of multiple protonation states of titratable groups within a peptide in response to a fixed environmental pH. This method has been used to investigate pH effects on peptides in many biologically-relevant cases, including illuminating the pH-dependence of the structures of the amyloid-*β* protein, a key peptide in the pathology of Alzheimer’s disease ^19^, and the influenza virus fusion process involving influenza fusion peptide insertion into the membrane of the host endosome ^20^.

In this article, we employ constant pH molecular dynamics (CpHMD) to examine the protonation states of GL13K lysine residues and their impact on the conformational dynamics of a single peptide. In Section 3.1, we investigate the *pK*_*a*_ values of the four lysine residues, to test the hypothesis that *different lysine residues in the GL13K sequence have different pK*_*a*_ *values* according to their differing local environments. Accepting this hypothesis would suggest a differential impact of the lysines in the sequence on the control of crystalline *β* sheet formation of GL13K at different pH levels and would support the idea of a single key lysine controlling aggregation. Next, in Section 3.2, we focus on the lysine residue located on the hydrophobic face (LYS11) (figure 1-b) and examine its protonation state. Specifically, we test the following hypothesis: *Residue 11 has a lower pK*_*a*_ *value than the other lysine residues*. A lower *pK*_*a*_ value would lead to deprotonation of the lysine on the hydrophobic face first and would support the hypothesis that it is the primary controlling factor of fibril formation. To our surprise, however, our results did not support this hypothesis, and thus we generated and tested a new hypothesis: *Residue 11 has a higher pK*_*a*_ *value than the other lysine residues*. To characterize the interplay between pH and accessible configurational space, we investigate the effects of pH level on *β* ladder content, end-to-end distances, and radius of gyration (Section 3.3). Finally, to better characterize the holistic interaction between pH and conformation, we explore metastable states using collective variable discovery, free energy landscape visualization, and Markov state models (MSM) and compare the conformational landscapes of isolated GL13K across different pH levels (Sections A.1, and 3.4).

## 2 Methods

### 2.1 Constant pH Molecular Dynamics Simulations

We conduct all-atom constant pH molecular dynamics (CpHMD) simulations of a single peptide in a water box using the *λ*-dynamics package for GROMACS version 2021^17,24,25^, ph-builder ^26^, with the Charmm36m force field modified for use with this package ^27^. We model the initial structure of GL13K using the PEP-FOLD4 web-server ^21^. We model the peptide with C-terminus amidation, meaning a neutral free carboxyl group, for consistency with previous simulations and experimental conditions ^7,28^. All lysine residues are modeled as titratable, while other residues are modeled as fixed-charge, including the N-terminal +1 charge, as none of the protonation states of the other residues are expected to change much, even at a pH of 12.5. We run five independent replicates for ten different pH levels, 8, 8.5, 9, 9.5, 10, 10.5, 11, 11.5, 12, and 12.5, for a total of 50 simulations. These pH levels were chosen to center on the *pK*_*a*_ of lysine in water (10.5). Although we note that simulating only a single peptide in aqueous solution will not permit us to investigate the more complex environment of multi-peptide aggregation, we argue that this investigation lays necessary high-precision theoretical groundwork for larger-scale understanding; additionally, we restrict ourselves to the experiments we planned and registered in advance on the Open Science Framework ([GL13K constant pH registration](https://doi.org/10.17605/OSF.IO/EW4VM) ^29^).

We place a single GL13K peptide in a cubic simulation box with an edge length of approximately 8 nm, ensuring a minimum padding of 2 nm from each face of the box such that the peptide does not see its periodic image. A steepest descent algorithm is used for energy minimization with a 0.01 step size and tolerance of 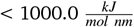 to a maximum of 5000 steps. Each replica undergoes 10 nanoseconds (ns) of standard NVT and NPT equilibration, followed by a 100 ns final production run under NPT conditions. For analysis purposes, the last 90 ns (post-equilibration) is used. The starting state of all titratable residues (lysine residues) is positively charged (*λ* =0). We model the water with the TIP3P water model ^30^, a 3-site rigid water model. In all simulations, we use physiological salt concentration (0.15 mM) of NaCl. The simulation temperature is set to 300 K. Electrostatic interactions are modeled with the particle mesh Ewald method ^31^ with a real-space cutoff of 1.2 nm and a 0.14 nm Fourier grid spacing, and van der Waals interactions as LJ potentials with a 1.2 nm cutoff. The v-rescale thermostat ^32^ is used with a time constant of 0.5 ps. The C-rescale barostat ^33^ is used to keep the pressure at 1 bar with a 5 ps period. Integration is performed with the leapfrog integrator ^34^ with a timestep of 2 fs. Bond length constraints to hydrogens are enforced with the LINCS algorithm ^35^ to fourth order and a single correction iteration. For sampling, we record the data every 0.02 ns, resulting in 4500 samples for each lysine residue per replica at each pH level. In total, for each lysine residue, we have 22500 samples over five replicates at each pH level.

#### 2.1.1 Average Deprotonated State and its relation to Henderson–Hasselbalch equation

In the *λ* -dynamics package for Gromacs introduced by Aho *et al*. ^17^, a non-terminal lysine residue has one titratable site (*ε*-amino group) attached to the furthest carbon (*ε*-carbon) from the *α*-carbon. In order to calculate the *pK*_*a*_ values of this titratable site for each lysine residue, we need to calculate the average fraction of the deprotonated state (*S*_*dep*_) across different replicas and various pH levels. The 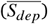 is calculated as

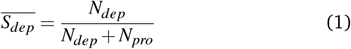

where *N*_*dep*_ and *N*_*pro*_ are the total number of frames in which the site is deprotonated and protonated, respectively, as determined from the *λ* trajectory. Following the recommendations of the package authors ^17,27^ we define the site as deprotonated if *λ* > 0.8 and protonated if *λ* < 0.2.

The relation between the pH of a solution, the *pK*_*a*_ of an acid, and the ratio of the concentrations of its conjugate base to the acid is described by the Henderson-Hasselbalch equation ^36^,

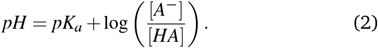

Since the ratio of concentrations can be assumed to be the same as the ratio of deprotonated to protonated frames, the fraction of the deprotonated state as a function of *pK*_*a*_ may be written,

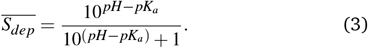

#### 2.1.2 Shape Analysis

To quantify GL13K conformations, we calculate a variety of shape parameters using the MDTraj scientific package ^37^. We compute the radius of gyration (*R*_*g*_), acylindricity (*a*_cyl_), and asphericity (*a*_sph_), which are all quantities derived from the principal moments 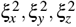 of the gyration tensor. Additionally, we calculate the end-to-end distance (*r*_*e*2*e*_) by finding the distance between *α* carbon of the first and the last residues.

Radius of gyration is defined as

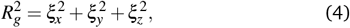

and is a measure of the overall compactness (low *R*_*g*_) or extendedness (high *R*_*g*_) of the protein.

Asphericity is calculated as

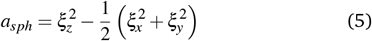

which shows how much the shape of a peptide deviates from a perfect sphere; a value of zero indicates a perfect sphere, while a value of one indicates a perfect line.

Acylindricity is calculated as

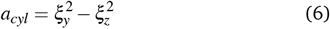

which shows how much the shape of a peptide deviates from a perfect cylinder; a value of zero indicates a perfect cylinder.

End-to-end distance is described as

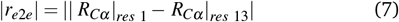

which measures the straight-line distance between the two terminal *C*_*α*_ atoms. Unlike the radius of gyration, this does not quantify the overall shape of the protein; a small *r*_*e*2*e*_ implies proximity of the termini but does not distinguish between a flat hairpin and a rounded loop.

#### 2.1.3 Secondary Structure Analysis

For further quantification of GL13K’s conformational tendencies, we assess the secondary structure of the peptide. We used the DSSP package from the MDTraj Python package ^38^ to identify the secondary structure of each residue at each timestep according to backbone angles and side chain positions. At each timestep *i*, the fraction *SSAF*_*S*_ of a residue with assigned type *S* was calculated as,

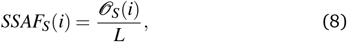

where 𝒪_*S*_(*i*), the occurrence of identifier *S* at the *i*th timestep, is the number of times an identifier *S* was found in timestep *i* and *L* = 13 is the length of the peptide. We investigate general secondary structure formation using the coarse-grained DSSP assignments of *S* ∈ {*H, E,C*} for *α*-helix, *β* sheet, and random coil, and we further investigate the different types of *β* sheet structures by using the more detailed version of DSSP, in which extended strand and isolated *β* -bridge fractions are defined by “E” and “B” identifiers.

#### 2.1.4. Markov State Model Construction and Analysis

A Markov State Model (MSM) is a computational method used to analyze and model the dynamics of complex systems, such as protein folding, to find slow processes that are interpreted as transitions between stable states. These stable states are long-lived (e.g. metastable) conformational states (i.e. free energy wells) where the peptide remains for a significant time before transitioning to another state by crossing a free energy barrier. The construction of an MSM assumes that one can represent the system with a model that possesses the Markov property, meaning the future state of an MSM depends only on its current state, not its past states. To construct a traditional MSM, there are six essential steps ^39^:

1. **Feature Selection**: Feature selection consists of identifying kinetically-relevant phase-space variables to describe the conformations of the system at each time step, such as peptide backbone dihedrals or distances between backbone atoms. Typically, such variables should be independent of system symmetries, i.e. rotation and translation.
2. **Dimensionality Reduction**: Dimensionality reduction, using methods like principal components analysis (PCA) or time-lagged independent component analysis (TICA), transforms high-dimensional features into a lower-dimensional space of collective variables (CVs) that is both easier to visualize and more robust for MSM construction while preserving important system dynamics. Thanks to collective motions, the dynamics of protein and peptide systems can be described by just a few dynamically or kinetically relevant variables ^40^.
3. **Discretization**: Discretization groups data described by CVs into kinetically relevant microstates using standard clustering techniques like k-means.
4. **MSM Estimation**: In the estimation step, one builds the transition probability matrix that describes the MSM between microstates based on the assumption of Markovianity.
5. **MSM Validation**: Via implied timescales and Chapman-Kolmogorov tests ^41^, one ensures that the assumption of Markovianity holds and that the model accurately captures the system kinetics.
6. **Coarse Graining the Microstates:** Coarse-graining microstates into macrostates using methods like Perron-Cluster Cluster Analysis (PCCA) or Robust Perron Cluster Analysis (PCCA+) ^42,43^ simplifies the MSM while retaining key dynamical properties for easier interpretation of metastable states.

Before constructing the MSM model for each pH level individually, we removed water and ions from the system and fixed the periodic boundary conditions (PBC) by making the peptide whole.

For dimensionality reduction, we used two different methods, Time-lagged Independent Component Analysis (TICA) ^44^ and principal component analysis (PCA) ^45^. PCA was used on all data simultaneously, to extract general principal components describing the overall conformational variance of the peptide as the pH varied. TICA CVs were determined separately for each pH level to better resolve specific kinetic details. As potential features for constructing the MSM, we considered proper and improper torsions, backbone atom distances, and backbone atom positions. Proper and improper torsions were selected for the input features to TICA, and only proper torsions were selected for the input features to PCA because they displayed the highest VAMP-2 score ^46^. To determine suitable lag times for dimensionality reduction with TICA at each different pH level, we investigated lag times ranging from 10 to 28 ns in increments of 2 ns (500 to 1400 steps, with each step being 20 ps). To choose the TICA lag time we used the following criteria: a) The VAMP2 score was > 1, showing the chosen feature has necessary information to distinguish between metastable states at that lag time, b) The 2D free energy surface built on top of the two independent components (IC1 and IC2) of TICA was clearly separated (clear states).

To determine suitable lag times for all MSMs, we used the following criteria: (a) the minimum lag time for which the implied timescales (ITS) plot demonstrates a plateau above the *ITs* = *τ* equivalency line to ensure Markovian behavior, as is standard in the literature ^47^ and (b) we confirm that the number of kinetic processes is in agreement with the number of stable states suggested by 2D FES. In figures S15, we plotted the implied timescales of the system at different pH levels.

For discretization, we used the k-means clustering method ^48^ to group the data based on similarities of features. For finding the optimal value for the number of clusters, we computed VAMP-2 scores across various cluster sizes and plotted them with confidence intervals. We chose the optimal number of clusters as the minimum number for which the plot of the VAMP-2 score displays a plateau. For MSM validation, we used the Chapman-Kolmogorov test ^41^. Finally, we grouped the microstates from previous steps into macrostates using the PCCA+ method ^42,43^, and we calculated the transition states and the mean first passage time (MFPT) between these states. We constructed individual MSM models in a TICA coordinate space employing the Deeptime package, version 0.4.5^49^ (Section 3.4). We constructed MSM models in a shared PCA coordinate space using the PyEMMA package, version 2.5.7^50^ (Supplementary Information.)

## 3 Results and Discussion

We performed constant pH molecular dynamics of GL13K at 10 different pH levels to understand the deprotonation states of GL13K and their effects on peptide conformational behaviour. We investigated the differences of the four lysine residue *pK*_*a*_ values, specifically, if the *pK*_*a*_ value of the last lysine residue is lower than that of the other lysine residues. In what follows, for each section, we explain our hypotheses and explicitly state whether the step was part of hypothesis testing or generation.

The analyses were conducted in accordance with our preregistered plan on the Open Science Framework (OSF; see [GL13K constant pH registration](https://osf.io/ew4vm/) ^29^). We tested the preregistered hypotheses as outlined in our OSF preregistration, while also exploring additional hypotheses generated during the study, which are discussed in the respective subsections below. The following hypotheses are part of hypothesis testing:

1. Different lysines in the GL13K sequence have different *pK*_*a*_ values.
2. Residue 11 (LYS11) has a lower *pK*_*a*_ than the other lysine residues.
3. Increasing pH will lead to loss of *β* strand content and decreasing radius of gyration and end-to-end radius.

The following hypothesis is part of hypothesis generation:

1. LYS11 has a higher *pK*_*a*_ value than the rest of the lysine residues.
2. Increasing pH leads to heightened *β* strand content
3. Increasing pH leads to subtle shifts in the free energy landscapes and stabilization of different kinetically-relevant metastable states

### 3.1 LYS11 has higher *pK*_*a*_ value than LYS5 but not LYS2 or LYS7

Initially, we start by expressing the first null and alternative hypotheses as outlined in the OSF plan as part of hypothesis testing:

- **Null Hypothesis (***H*_0_): Different lysine residues in the GL13K sequence possess equal *pK*_*a*_ values.

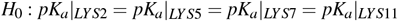
- **Alternative Hypothesis (***H*_1_): Different lysines in the GL13K sequence possess different pKa values.

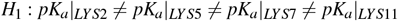

For rejecting or accepting the null hypothesis, we calculate the deprotonation fraction for each lysine residue at each pH level across 5 replicas according to equation 1. The *pK*_*a*_ is extracted for each lysine by fitting the Henderson-Hasselbalch equation to the average deprotonation at each pH level according to equation 3. Additionally, to quantify the precision and uncertainty of our extracted *pK*_*a*_ values, we estimate the variability by extracting the maximum and minimum possible *pK*_*a*_ values given the deprotonation data for each replica. If the *pK*_*a*_ values of different residues do not overlap to within these ranges of uncertainty, we interpret it as the *pK*_*a*_ values differing. Additionally, we quantify the magnitude of the difference as a measure of the effect size.

Figure 2 shows the calculated *pK*_*a*_ values and uncertainty level for each lysine residue (mean *pK*_*a*_ values for LYS2, LYS5, LYS7, and LYS11 are 10.211, 10.117, 10.194, and 10.305, respectively. The ranges are LYS2:[10.157,10.237], LYS5:[9.968,10.186], LYS7:[10.088,10.248], and LYS11:[10.233,10.355]). Except for LYS5 and LYS11, the rest have a mutual overlapping of *pK*_*a*_ values, so we reject the null hypothesis for the case of a single GL13K peptide in water. For LYS5 and LYS11, we observe a clear separation in uncertainty levels, indicating statistical significance; however, the small difference (≈ 0.0466) with percent difference of average *pK*_*a*_ values between LYS5 and LYS11 defined as

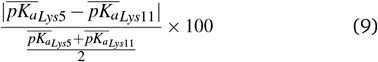

is 1.84 %, which suggests a small effect size.

**Fig. 2.**
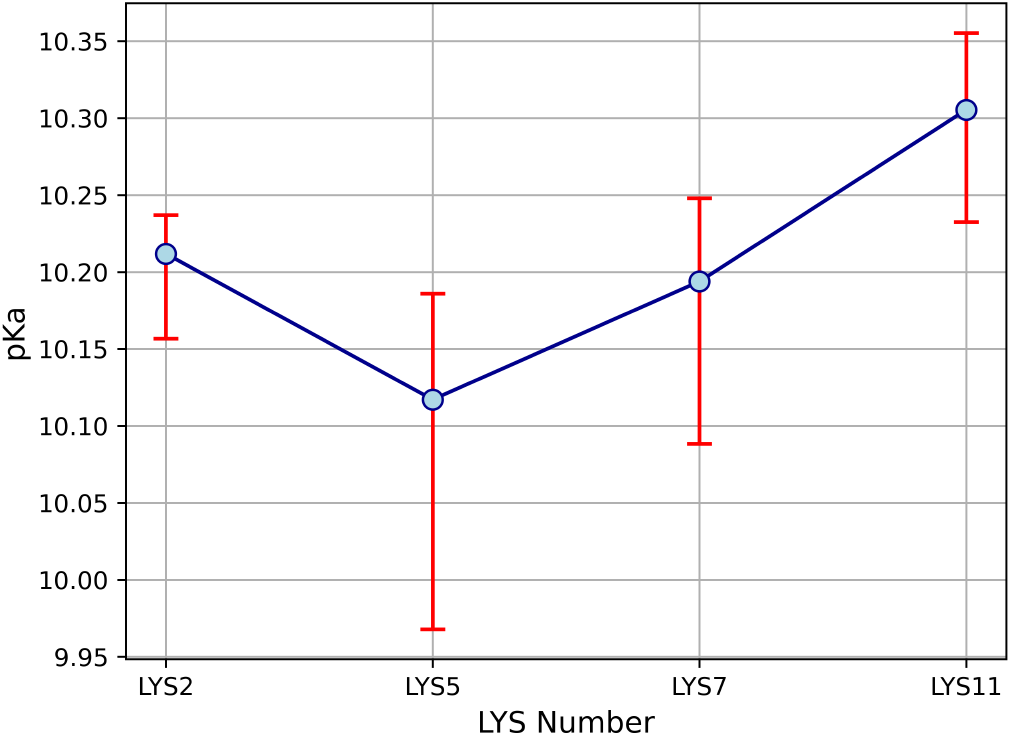
The *pK*_*a*_ values and uncertainty level of all four lysine residues calculated by fitting the Henderson-Hasselbalch equation to the average deprotonation at each pH level.

### 3.2 LYS11 has a significant deprotonation ratio difference with other lysine residues at lysine side chain *pK*_*a*_ value

As previously mentioned, GL13K does not self-assemble into fibrils at pH 7.4 but does form fibrils at higher pH values. A previously advanced hypothesis for this behavior is that it is controlled by the *pK*_*a*_ value of the lysine residue on the hydrophobic face (LYS11). Therefore, in this section, we focus on this lysine, near the C terminus, employing the next hypothesis:

- **Null Hypothesis (***H*_0_): LYS11 located on the hydrophobic face has the same *pK*_*a*_ value as other lysine residues.
- **Alternative Hypothesis (***H*_1_): LYS11 located on the hydrophobic face has a lower *pK*_*a*_ value than other lysine residues.

In other words, we investigate whether LYS11 has a lower *pK*_*a*_ value than the rest of the lysine residues (hypothesis testing). To test this hypothesis, at each pH level, we quantify the number of frames where LYS11 is protonated/deprotonated and compare it to the number of frames where other lysine residues are protonated/deprotonated. To avoid sampling non-physical states, a biasing potential term in the potential energy function ensures that the majority of *λ* values lie close to *λ* = 0 or *λ* = 1^17^. Therefore, after sampling the *λ* trajectory at each pH level, the resulting distributions are either unimodal (for primarily deprotonated or primarily protonated systems) or bimodal (for systems displaying both protonated and deprotonated character over the course of the simulation). Figure 3 shows these sampled distributions for all four lysine residues at each pH level. To rigorously distinguish the unimodal from the multimodal (bimodal) distributions, we assess the bimodality of each lysine’s *λ* -value data set at each pH level using Hartigans’ dip test ^51^, with a significance threshold of *p* < 0.05. In our study, a significant result indicates a bimodal distribution. The *λ* values of pH 8 and 12.5 show unimodal behavior for all lysines, and the rest passed Hartigans’ dip test (*p* values reported in Fig. 3 and Table S1). For systems that passed Hartigans’ dip test, the bimodal systems, we apply a Gaussian mixture model ^52^ to characterize the protonated and deprotonated states, yielding a Gaussian distribution per mode. Figure S1 shows the assigned Gaussian distribution functions of each lysine residue per pH level. In the next step, we assess the data set normality using the Quantile to Quantile (QQ) plot ^53^ and the Shapiro-Wilk test ^54^. Figures S2, S3, and S4 show the QQ plot for the data set from the first mode, the second mode of bimodal distribution functions, and unimodal distribution functions, revealing a non-normally distributed function for all these datasets. Additionally, in Table S2, p-values of the Shapiro-Wilk test are presented, demonstrating non-normally distributed functions. Moreover, we assess the variance equality of data sets for all the possible comparisons between LYS11 and other lysine residues using the Levene test ^55^. In Table S3 we present the p-value (p-value threshold is set at 0.05) regarding the Levene test, showing that the majority of comparisons do not have variance equality. At this point, we demonstrated that the data sets do not follow a normal distribution function and do not meet the variance equality condition. Therefore, for comparison between LYS11 deprotonation ratio and other lysine residues, we perform a non-parametric test, the Mann-Whitney U test ^56^, on the data sets.

**Fig. 3.**
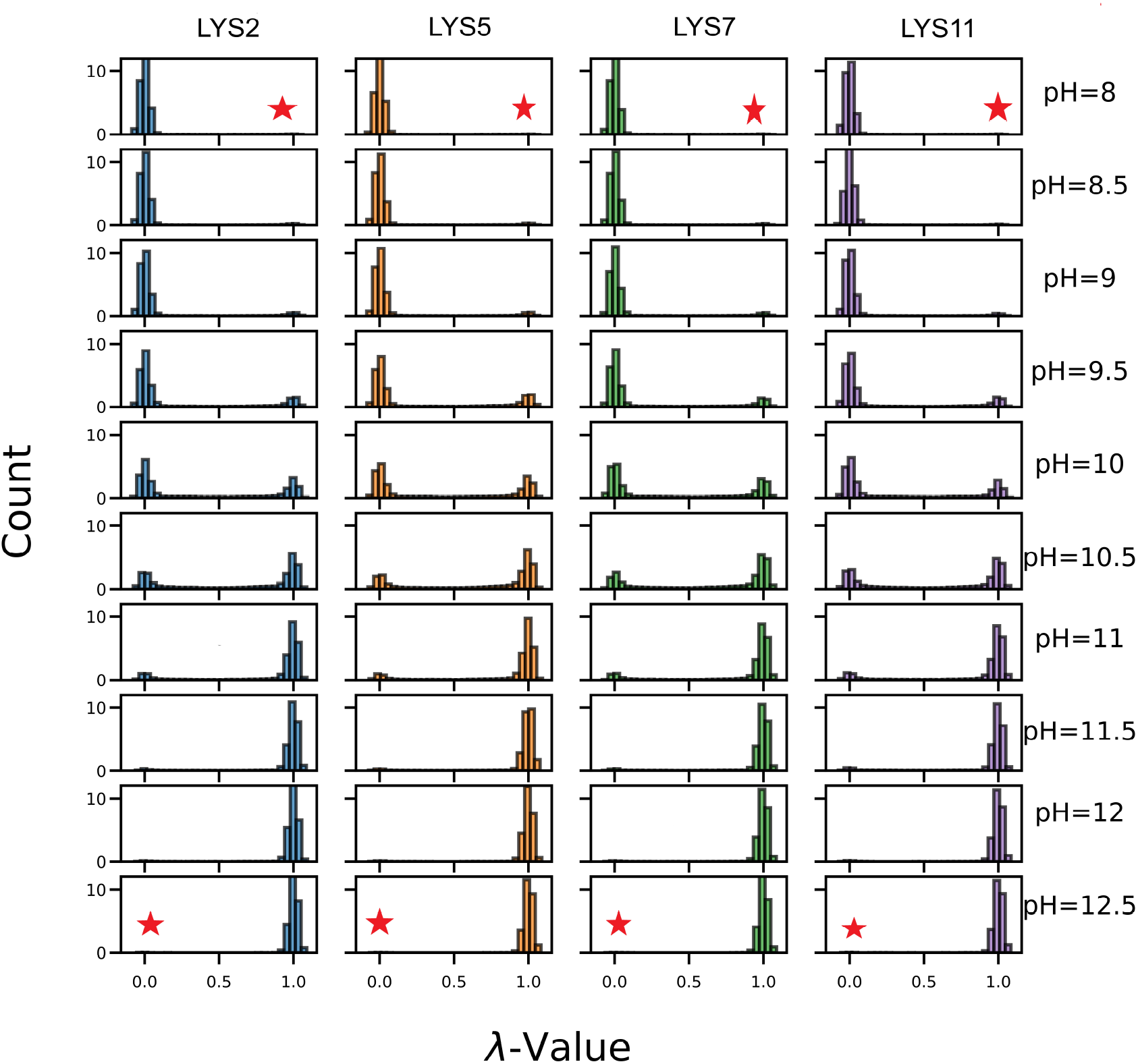
*λ* values of 4 lysine residues at 10 different pH levels. The significance threshold of *p* < 0.05 is set to distinguish unimodal from bimodal distribution functions. The unimodal distribution functions are marked with a small red asterisk.

To assess whether we can reject the null hypothesis that LYS11 is protonated at the same rate as each of the other lysine residues at each of the 10 pH levels, we perform a number of one-sided Mann-Whitney U tests using an asymptotic method (large sample size) with Bonferroni correction ^57^. For the comparison, we have 4 lysine residues and 10 different pH levels, resulting in 30 comparisons. Aiming at a traditional familywise error rate of 0.05, we set the significance level for each test at 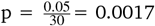. Table 1 presents the one-sided Mann-Whitney p-values for 30 comparisons, but these p-values are consistently large (close to 1), indicating no significant differences between the groups. In addition, we calculated the titration curve of each lysine residue at all different pH levels by fitting the deprotonation ratio into the Henderson-Hasselbach equation described in the methods section (equation 3) and plotting the average *pK*_*a*_ value.

**Table 1.** *p*-Values from one-sided Mann-Whitney Test with initial hypothesis for hypothesis testing (“LYS11 has a lower *pK*_*a*_ value than the other lysines.”). Significance level set at 0.0017 for a family-wise error rate of 0.05. We highlight in bold *p*-values for the test that lie beneath the significance level.

|  | pH=8 | pH=8.5 | pH=9 | pH=9.5 | pH=10 | pH=10.5 | pH=11 | pH=11.5 | pH=12 | pH=12.5 |
| --- | --- | --- | --- | --- | --- | --- | --- | --- | --- | --- |
| LYS2 to LYS11 | 9E-1 | 1 | 1 | 7E-1 | 1 | 1 | 7E-1 | 1 | 1 | 9E-1 |
| LYS5 to LYS11 | 9E-1 | 1 | 1 | 1 | 1 | 1 | 1 | 1 | 7E-1 | 2E-1 |
| LYS7 to LYS11 | 8E-1 | 1 | 1 | **1E-4** | 1 | 1 | 1 | 1 | 8E-1 | 3E-1 |

Additionally, we plotted the average deprotonation ratio for each pH level across all replicas in Figure 4. The error bar is calculated as the standard error of independent replicas. In figure S5, we plotted the titration curve of each lysine residue for each replica. From both Figs 2 and 4 we observe that LYS11 not only lacks a lower *pK*_*a*_ value compared to other lysine residues but also exhibits a shift in the other direction, suggesting a potentially higher *pK*_*a*_ value instead, concomitant with especially at pH levels near the *pK*_*a*_ of lysine in water. From these results, we generate a new alternative hypothesis as such: “*LYS11 has a higher pK*_*a*_ *value than the rest of the lysine residues*” (hypothesis generation). For assessing this hypothesis, we perform the same statistical test with the correct one-sided alternatives. The p-values are reported in Table 2. The results show a significant difference between LYS11 and LYS5 deprotonation ratios at pH levels of 8.5 to 11.5. Generally, LYS11 has a significantly different deprotonation ratio at pH 10, pH 10.5, and pH 11.5 from other lysine residues, specifically at the reported experimental *pK*_*a*_ value of lysine residue ^58^. Thus, we reject the null hypothesis in favor of the newly-generated alternative hypothesis that LYS11 has a *higher pK*_*a*_ value than other residues as measured by the distributions of *λ* values.

**Table 2.** *p*-Values from one-sided Mann-Whitney Test with modified hypothesis from hypothesis generation (“LYS11 has a higher *pK*_*a*_ value than the other lysines.”) Significance level set at 0.0017 for a family-wise error rate of 0.05. We highlight in bold *p*-values for the test that lie beneath the significance level.

|  | pH=8 | pH=8.5 | pH=9 | pH=9.5 | pH=10 | pH=10.5 | pH=11 | pH=11.5 | pH=12 | pH=12.5 |
| --- | --- | --- | --- | --- | --- | --- | --- | --- | --- | --- |
| LYS2 to LYS11 | 7E-2 | **1E-5** | **8E-9** | 2E-1 | **7E-33** | **4E-27** | 3E-1 | **4E-12** | 1E-2 | 1E-1 |
| LYS5 to LYS11 | 8E-2 | **2E-13** | **5E-12** | **5E-43** | **8E-70** | **1E-104** | **8E-5** | **2E-10** | 3E-1 | 8E-1 |
| LYS7 to LYS11 | 2E-1 | **2E-4** | **4E-5** | 9E-1 | **2E-43** | **1E-61** | 1E-2 | **8E-9** | 2E-1 | 7E-1 |

**Fig. 4.**
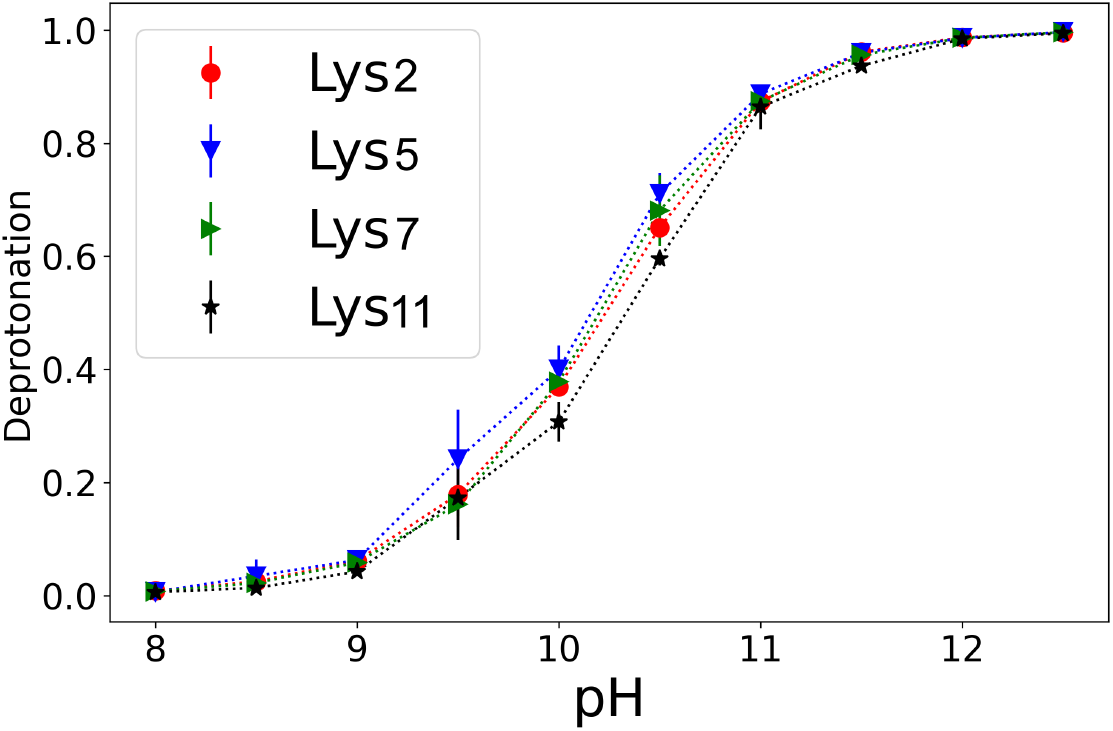
Titration curves of the lysine residues as titratable sites in GL13K peptide. The error bars indicate the standard error in the means of the five different replicas. Connecting lines are included to guide the eye and do not represent fits.

### 3.3 Increasing pH stabilizes collapsed conformations partly through the stabilization of transient *β* strands

Another important aspect of peptides is their conformational shape, including radius of gyration (*R*_*g*_), end-to-end distance (*e*2*e*), acylindricity (*a*_*cyl*_ ), asphericity (*a*_*sph*_), and secondary structure. In this section, we focus on the effects of pH on peptide conformation. Due to lysine residues charge-charge interactions, the peptide may not be able to form a more compact conformation. However, by increasing the pH and crossing the *pK*_*a*_ values of lysine residues, these charges are neutralized, and more compact structures may be seen. We tested the hypothesis that increasing pH would lead to decreasing radius of gyration and end-to-end radius, as well as loss of *β* strand content. We found support for the decrease of radius of gyration and end-to-end radius, but we found an increase rather than a decrease of *β* strand content, as tested by the correlation coefficients between pH level and the relevant parameters (Table 3).

**Table 3.** Pearson correlation coefficient between pH level and shape parameters, and between pH level and secondary structure propensity.

| Variable | $R_g$ | e2e | $a_{sph}$ | $a_{cyl}$ | Strand Fraction |
| --- | --- | --- | --- | --- | --- |
| pH | -0.48 | -0.42 | -0.49 | -0.23 | 0.32 |

**Table 4.** Lag time (steps) for constructing TICA and MSM models at each pH level.

|  | pH=8 | pH=8.5 | pH=9 | pH=9.5 | pH=10 | pH=10.5 | pH=11 | pH=11.5 | pH=12 | pH=12.5 |
| --- | --- | --- | --- | --- | --- | --- | --- | --- | --- | --- |
| TICA | 1100 | 1100 | 1200 | 1400 | 1100 | 1400 | 1400 | 1000 | 1000 | 1100 |
| MSM | 600 | 400 | 300 | 600 | 1100 | 1000 | 1100 | 1300 | 1000 | 1300 |

Figure 5 shows average values of *R*_*g*_, *r*_*e*2*e*_, *a*_*cyl*_ and *a*_*sph*_ at 10 pH levels. The standard error for each pH level is calculated across five independent replicates and indicated as the error bar. We observe moderate change in the shape parameters with pH. There is a tendency towards lower radius of gyration and higher variance above pH 10 (cf. also Fig. S6) concomitant with a tendency towards lower end to end distance and asphericity and higher variance in both (cf. also Figs S7, S8), but almost no change in acylindricity (cf. also Fig. S9). To quantify the effect size of pH increment on peptide shape descriptors, we calculate the Pearson correlation coefficient between pH and each one of these descriptors. The Pearson correlation coefficient measures the strength and direction of a linear relationship between two variables ^59^. In Table 3, we present the Pearson correlation coefficient between pH and peptide shape descriptor variables including *R*_*g*_, *e*2*e, a*_*cyl*_, *a*_*sph*_, and *β* -strand fraction. As expected, we observe a moderate negative linear relation between pH and *R*_*g*_, *e*2*e*, and *a*_*sph*_ and a weak negative linear relation between pH and *a*_*cyl*_ .

**Fig. 5.**
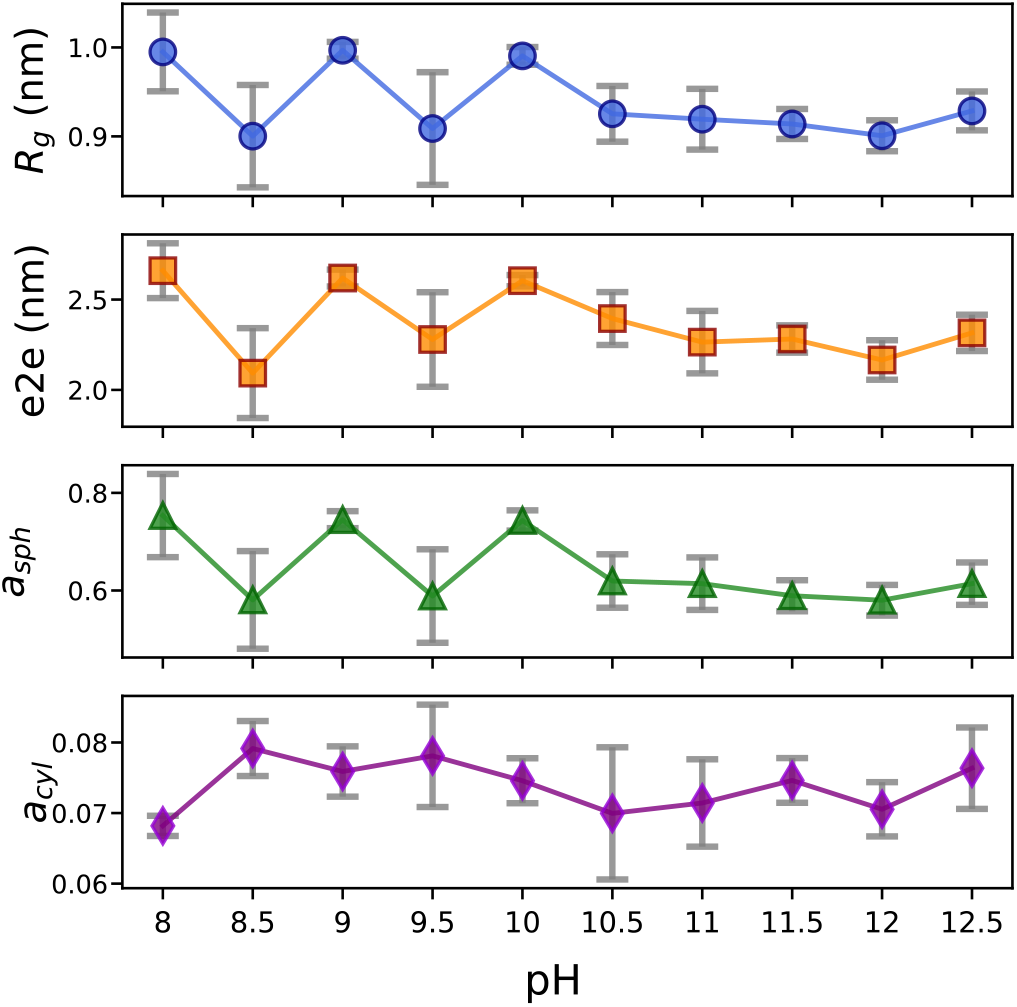
Average shape parameters calculated for 10 pH levels from top to bottom: radius of gyration (*R*_*g*_), end-to-end distance (*e*2*e*), asphericity (*a*_*sph*_), and acylindricity (*a*_*cyl*_). The error bars indicate the standard error in the means of the five different replicas.

Taken together, this indicates that for environmental pH values > 10 when significant deprotonation events begin to be observed (cf. Fig. 3), more compact conformations are moderately stabilized; whereas below these pH values, both extended and compact conformations are accessible, with a preference for extended conformations. In addition, compact conformations become more stable as the pH value of the environment increases.

Figure 6 shows the average secondary structure of the peptide over different pH levels, focusing on isolated *β* -bridge and extended strand percentage, with the standard error computed across five independent replicates. In a single peptide, an isolated *β* -bridge is a minimal *β* -sheet-like structure, defined by a single hydrogen bond between two residues, creating a localized interaction. On the other hand, an extended strand is a broader *β* -sheet-like conformation, involving a longer segment of the peptide and potentially the formation of multiple hydrogen bonds between residues. In this figure, all systems across various pH levels exhibit approximately between 90% to 95% coil structure, aligning closely with prior fixed-charge computational studies on the isolated GL13K peptide in solution ^7^, although the earlier report indicated a slightly higher percentage of helical and *β* -strand structures in uncharged GL13K. It is also important to note that the majority of helical structures are formed at a pH level below 10, where all four lysines are primarily found in the protonated (charged) state (cf. Fig. 3, top four rows), but *β* -strand structures are exclusively observed at pH levels of 10.5 and above, where all four lysines are found substantially or exclusively in the deprotonated (uncharged) state (cf. Fig. 3, bottom five rows). Indeed, we observe a moderate positive linear relation between pH and the strand fraction of secondary structure (Table 3). (At pH 10, we observe almost entirely random-coil conformations.) It is of note that, at the reported experimental *pK*_*a*_ value of the lysine side chain and close to our calculated *pK*_*a*_ for all LYS residues (10.5), we observed the emergence of *β* -sheet structures for the first time. This suggests a critical role of lysine’s protonation state in stabilizing these secondary structural elements. In figures S10, S11, and S12, we plotted the average helical, strand, and coil-like fraction per residue at each pH level, respectively. Specifically, in the figure S11, we observed that the *β* -sheet contents are mainly formed between LYS7 and LYS11, suggesting the important role of the partial deprotonation of those two residues in stabilizing intra-peptide *β* -sheet contents.

**Fig. 6.**
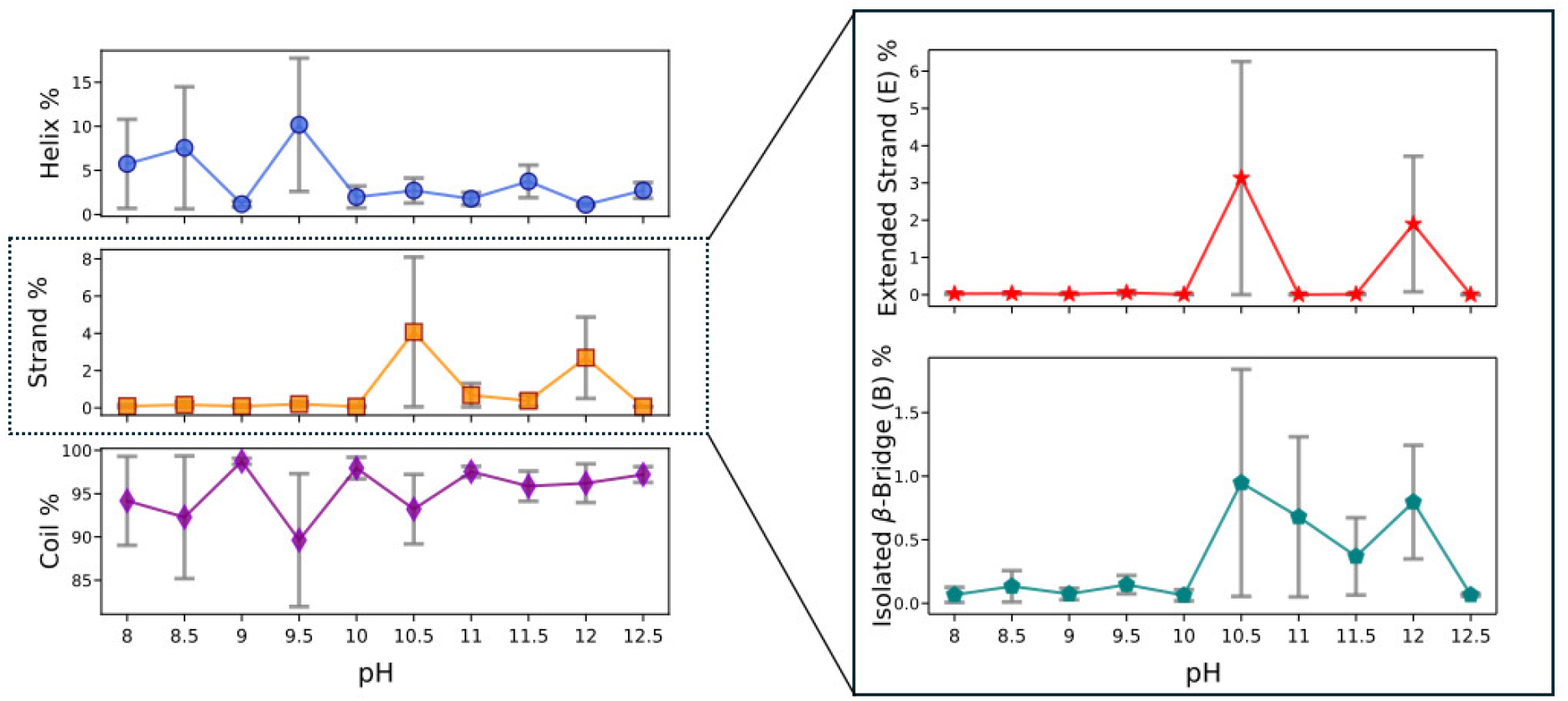
Secondary structure of peptide at each pH level is calculated. (left) Three categories of secondary structure for each pH level are plotted with standard error as error bars. (right) Two components of the Strand structure, isolated *β* -bridge and extended strand, are plotted for each pH level.

Based on the results, we argue that we can reject the null hypothesis. However, the Pearson correlation coefficient between pH and strand fraction is approximately 0.3, indicating a weak or moderate effect size. It is important to mention that a vast number of AMPs are disordered in a non-active environment, such as solution ^60^.

### 3.4 Changing pH subtly modifies accessible conformational states and highlights *β* hairpin configuration near the *pK*_*a*_ of lysine

Free energy surfaces (FES) can reveal metastable states, barriers between these states, and frequency of visited states of the system. The change in Helmholtz free energy with respect to a given reaction coordinate *χ* (Δ*F*(*χ*)) can be calculated as

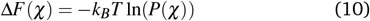

where *k*_*B*_ and T are the Boltzmann constant and temperature, respectively, and *P*(*χ*) is the probability of obtaining a particular reaction coordinate value, which may be estimated through binning across trajectories. Highly-populated states possess correspondingly lower free energies. Markov State Models can be used to estimate transition rates between such metastable states.

In this article, we construct and compare free energy surfaces in three different sets of reaction coordinates for three different and complementary purposes as part of hypothesis generation. In Fig. 7, we display the free energy landscapes at different pH levels in shared collective variables to identify whether there are major differences in occupancy of the overall free energy landscape. In Fig. 7-a, we employ the top two PCs from a PCA projection of the backbone and sidechain torsions, employing all configurations from all simulations at each pH level to identify high-variance collective motions sampled by all peptides. In Fig. 7-b, we employ the first and second moments of the gyration tensor to compare the free energy of simple interpretable collective motions known to well-describe short peptides. In Fig. 8-a, we visualize separate TICA projections for each pH level. Fig. 8-b, we depict the same landscape colored by metastable states obtained from individually-fit MSMs, with mean first passage times (MF-PTs) indicated for transitions between states.

**Fig. 7.**
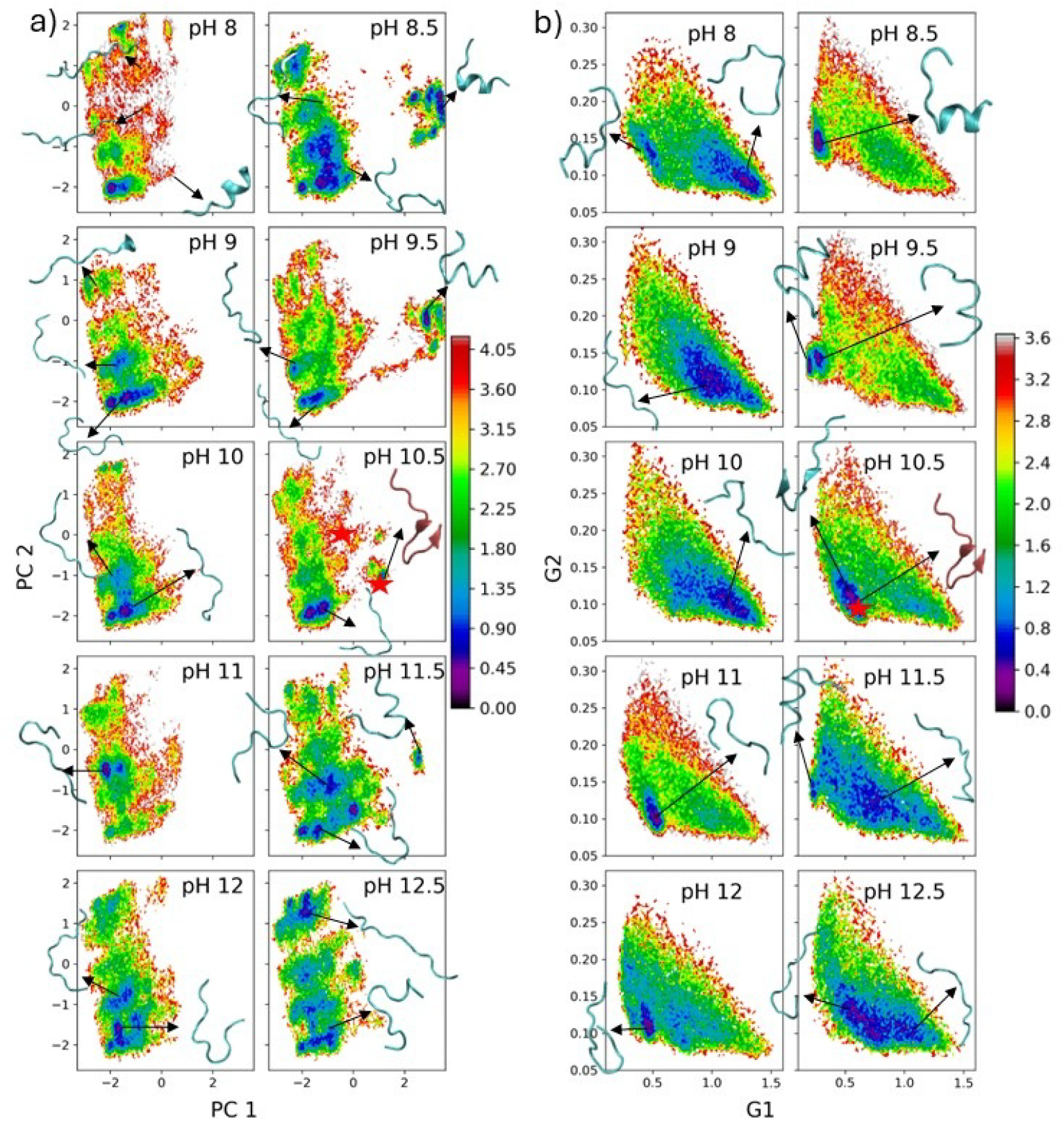
Two-dimensional free energy surfaces of a single GL13K in solution at different pH levels, (a) applying PCA transformation obtained from all the data across 10 pH levels on individual pH levels. (b) Plotting simple FES using 1st and 2nd moments of the gyration tensor. The red asterisk represents the location of the short anti-parallel beta sheet in these methods.

**Fig. 8.**
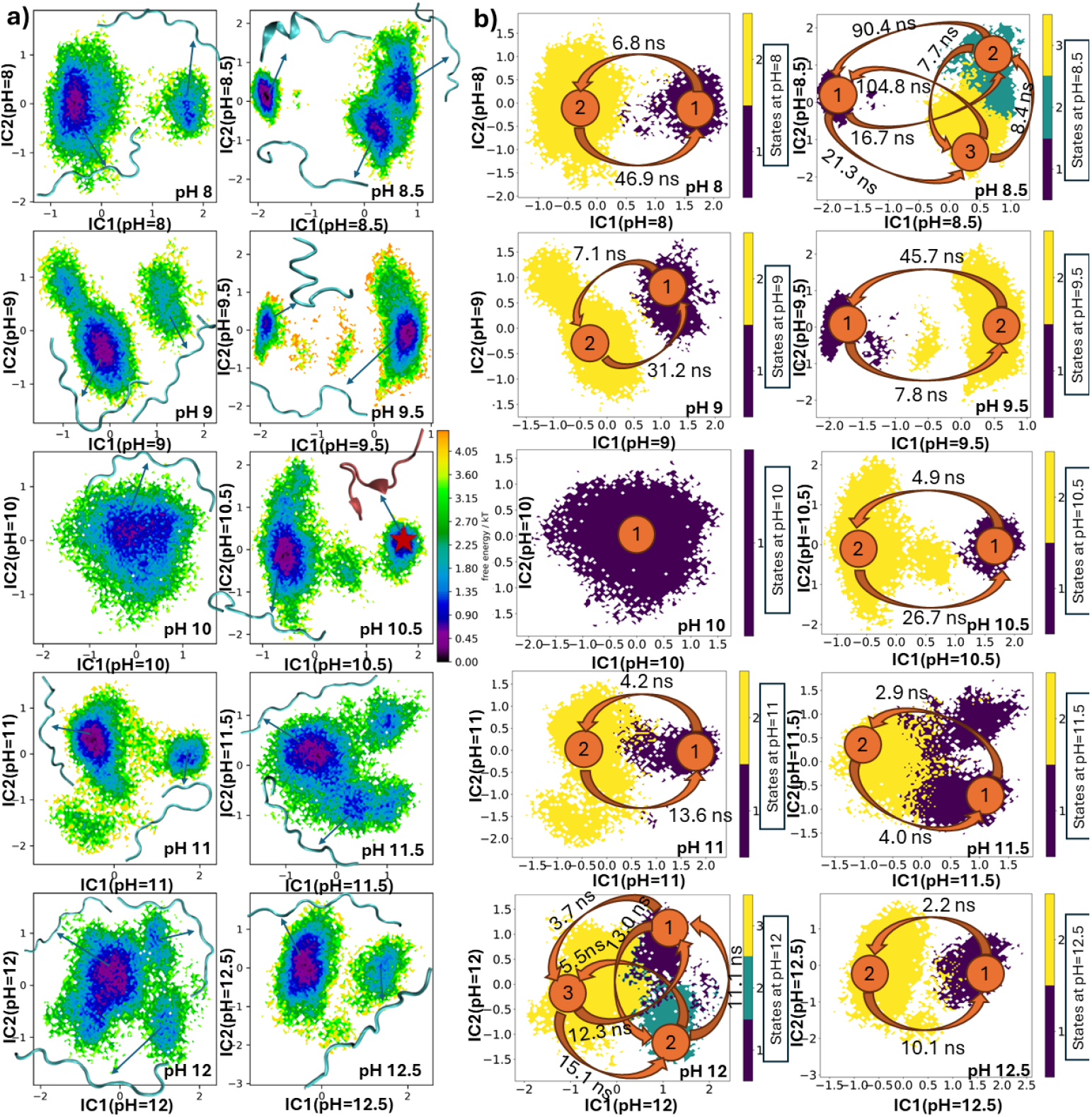
Free Energy Surface (FES) of the system constructed by the first two independent components (ICs) at all 10 pH levels computed using TICA as the dimensionality reduction method, colored by (a) free energy values and (b) metastable states, with mean first passage times (MFPTs) indicated for transitions between states.

In all figures the differences in the landscapes are quite small. In particular, in Fig. 7b, the overall shapes of the peptides as described by the moments of the gyration tensor are very similar. Key points of note that rise to about the order of thermal fluctuations (∼ 5*k*_*B*_*T* ) in the occupancy of the overall free energy landscape: As shown in Fig. 7a are that the upper left corner (low PC1 values, high PC2 values) appears to correspond to more extended conformations, while the generally well-populated regions of the lower left corner (*PC*1, *PC*2 < 0) appear to correspond to unstructured collapsed or semi-collapsed states. We observe more structured conformations (helix, near-helix, *β* hairpin) for the right-hand side of the free energy surface plot (*PC*1 > 0). In particular, at pH 8.5 and 9.5 we see a folded-over helix-like configuration in a region near *PC*1 ≈ 3, which is only populated at these two pH levels. At pH 10.5 we also observe a *β* hairpin configuration at around *PC*1 = 1, *PC*2 = −1, a region that is only populated for *pH* > 8.5 (when there is at least some partial deprotonation). Comparing to the MSMs built on the individual pH levels, we do observe for nearly all levels 2-3 states separated by shallow free energy barriers with transition times generally on the order of about 10 ns (the exception being at pH 10, where only a single state is resolved). However, we see that at pH 8.5 and 9.5, the folded-over helix-like configuration is resolved as a state, with a transition time into it of about 50-100 ns from the other states. Additionally, at pH 10.5, the *β* hairpin configuration is once more resolved as a separate state, confirming its stability and appearance primarily near the *pK*_*a*_ of the lysine residues.

We note that the comparatively small scale of the differences in the free energy surfaces and the comparatively rapid transition times underscore that the effects of pH represent subtle shifts in an overall disordered landscape, which is expected given the relative scarcity of secondary structure observed and the short length of the peptide.

## 4 Conclusions

We performed constant-pH molecular dynamics simulations of a single GL13K antimicrobial peptide in solution at 10 different pH levels from 8 to 12.5. We investigated the *pK*_*a*_ values of 4 lysine residues with the null hypothesis that different lysine residues in the GL13K sequence have the same *pK*_*a*_ values. Although we did not reject this null nypothesis, we did demonstrate that we were able to reject the null hypothesis that LYS11 has the same *pK*_*a*_ value as the others, and we measured the *pK*_*a*_ of the lysines in the sequence to be lower than the experimental *pK*_*a*_ of lysine alone in aqueous solution. We identified a small but significant effect in that LYS11 is less deprotonated than the other lysines in the sequence at the same pH level. These observations have several implications. Firstly, except for pH levels around 10-11, we confirm that it is appropriate to model all lysines as protonated (*pH* < 10) or all lysines as deprotonated (*pH* > 11), which is important for simulation of larger scale aggregation at pH levels distant from 10.5. Secondly, from previous work we know that in formation of small-scale aggregates, *β* sheet content appears on the N-terminal side of the peptide, including LYS2, LYS5, and LYS7 when the peptides go from charged to uncharged, whereas a small amount of *β* sheet content can be found on the C-terminal side, including LYS11, regardless of charge state ^7^. Between this evidence and the evidence that LYS 11 has a slightly higher *pK*_*a*_ than the other three lysines, this argues that rather than LYS11 controlling aggregation, aggregation is primarily controlled by simultaneous deprotonation of LYS2, LYS5, and LYS7.

In addition to studying the *pK*_*a*_ values of the different lysines, we also probed the effect of pH level directly on the conformational states of the single-peptide system. We observed that increasing the pH results in a decrease of *R*_*g*_ and *r*_*e*2*e*_ corresponding to increasing collapse of the peptide due to removing the lysine charge-charge repulsion. We obtained a more detailed view of the changing conformationally-accessible ensemble by constructing free energy surfaces and Markov State models. Although the population of shared free energy surfaces was very similar and for all pH levels there is an expected preponderance of random coil configurations in solution for dilute or single peptides, we found subtle differences in the resolved metastable states when projecting into the most kinetically-relevant variables at each pH level.

When the pH is increased, we observed a transition from stabilization of *α* helical conformations to stabilization of a *β* -hairpin structure near the *pK*_*a*_ of the lysine residues to more random-coil conformations at higher pH. This suggests that for applications in which the *β* hairpin configuration plays a strong role, fixed-charge simulations may be insufficient, particularly due to the likely role of the protonation state of LYS11 to the stabilization of this hairpin. Notably, the observed elusive *β* hairpin configuration presents at the top of the hook the residue SER9, which is known to play a very significant role in lipopolysaccharide-GL13K binding ^5^. Indeed, mutation of the ninth residue away from serine significantly decreases GL13K binding to LPS, removing its potential protective effect against this bacterial endotoxin. Based on the results in the current article, we propose the following experimentally-testable hypothesis: that LPS binding is strongest at a pH level between 10 and 11. Should this hypothesis be validated, it would provide important evidence for the relevance of the configuration in question; should it be disproved, it would direct research towards other potential mechanisms of SER-controlled binding. Either of these outcomes would be valuable for optimizing the LPS binding of GL13K or for designing new peptides with heightened affinity for LPS. In future, this might be a path towards design of peptides that protect against bacterial virulence without incurring a resistance penalty (*i*.*e*. if the peptide neutralizes the toxin without killing the bacteria).

We note that our work is limited to dilute peptide conditions and further work is needed to address the question of peptide-membrane interaction or peptide aggregation near pH 10-11. An important open question remaining in the field is the local pH of a bacterial membrane, which is difficult to measure experimentally or model computationally. We have now pinpointed the pH range for which we can be relatively sure that fixed-charge modeling is sufficient. Another important area of future work will be modeling LPS-GL13K interactions at constant pH near and distant from the identified *pK*_*a*_ of LYS11, which would complement the experimental studies proposed in the previous paragraph.

## Supporting Information

Supporting information is available, including 3 tables and 19 figures.

The datasets containing molecular dynamics trajectories without water, structures, force fields, molecular dynamics input parameters, and analysis files are publicly available at the following DOI:*https://doi.org/10.5281/zenodo.17240858*. Full trajectories, including the positions of water molecules, are available from the authors upon reasonable request.

## Acknowledgment

This research was supported in part by Discovery Grant #RGPIN-2021-03470 from the National Sciences and Engineering Research Council of Canada. This research was enabled in part by support provided by Calcul Quebec (www.calculquebec.ca) and the Digital Research Alliance of Canada (https://alliancecan.ca). This research was undertaken, in part, thanks to funding from the Canada Research Chairs Program under grant number CRC-2020-00225.

## A Supporting Information

Figures S10, S11, and S12 show the average secondary structure and uncertainty level of peptide per residue for helix, strand, and coil fraction over different pH levels, respectively.

### A.1 High pH results in dominance of coil structure stable states

Understanding stable states of an antimicrobial peptide reveals the structural behavior of that peptide in the environment. In Figures S18, and S19, we built Markov state models using PCA transformation fitted on all the data across 10 pH levels in figures S18, and S19. We observed that PCA is not able to separate stable states very well, especially in pH levels of 12 and 12.5.

**Fig. S1.**
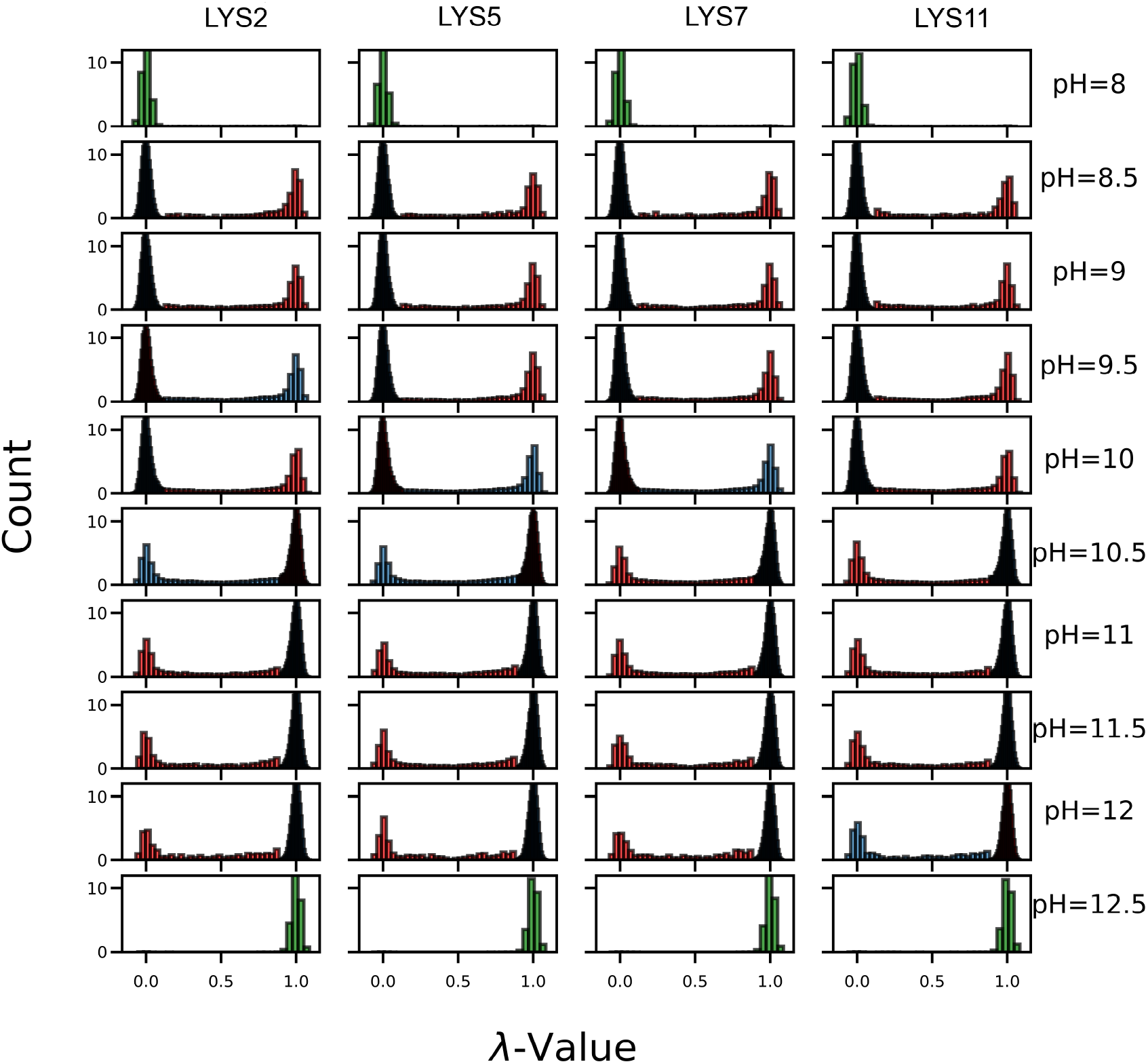
Assigned Gaussian distribution functions of each lysine residue per pH level. In each panel, different color represents the data assigned to a Gaussian distribution function.

**Table S1.** P-Values from Hartigan’s Dip Test.

|  | pH=8 | pH=8.5 | pH=9 | pH=9.5 | pH=10 | pH=10.5 | pH=11 | pH=11.5 | pH=12 | pH=12.5 |
| --- | --- | --- | --- | --- | --- | --- | --- | --- | --- | --- |
| LYS2 | 4E-1 | 0 | 0 | 0 | 0 | 0 | 0 | 0 | 4E-2 | 9E-1 |
| LYS5 | 7E-1 | 0 | 0 | 0 | 0 | 0 | 0 | 0 | 1E-2 | 9E-1 |
| LYS7 | 6E-1 | 0 | 0 | 0 | 0 | 0 | 0 | 0 | 3E-2 | 7E-1 |
| LYS11 | 9E-1 | 3E-3 | 0 | 0 | 0 | 0 | 0 | 0 | 9E-4 | 8E-1 |

**Fig. S2.**
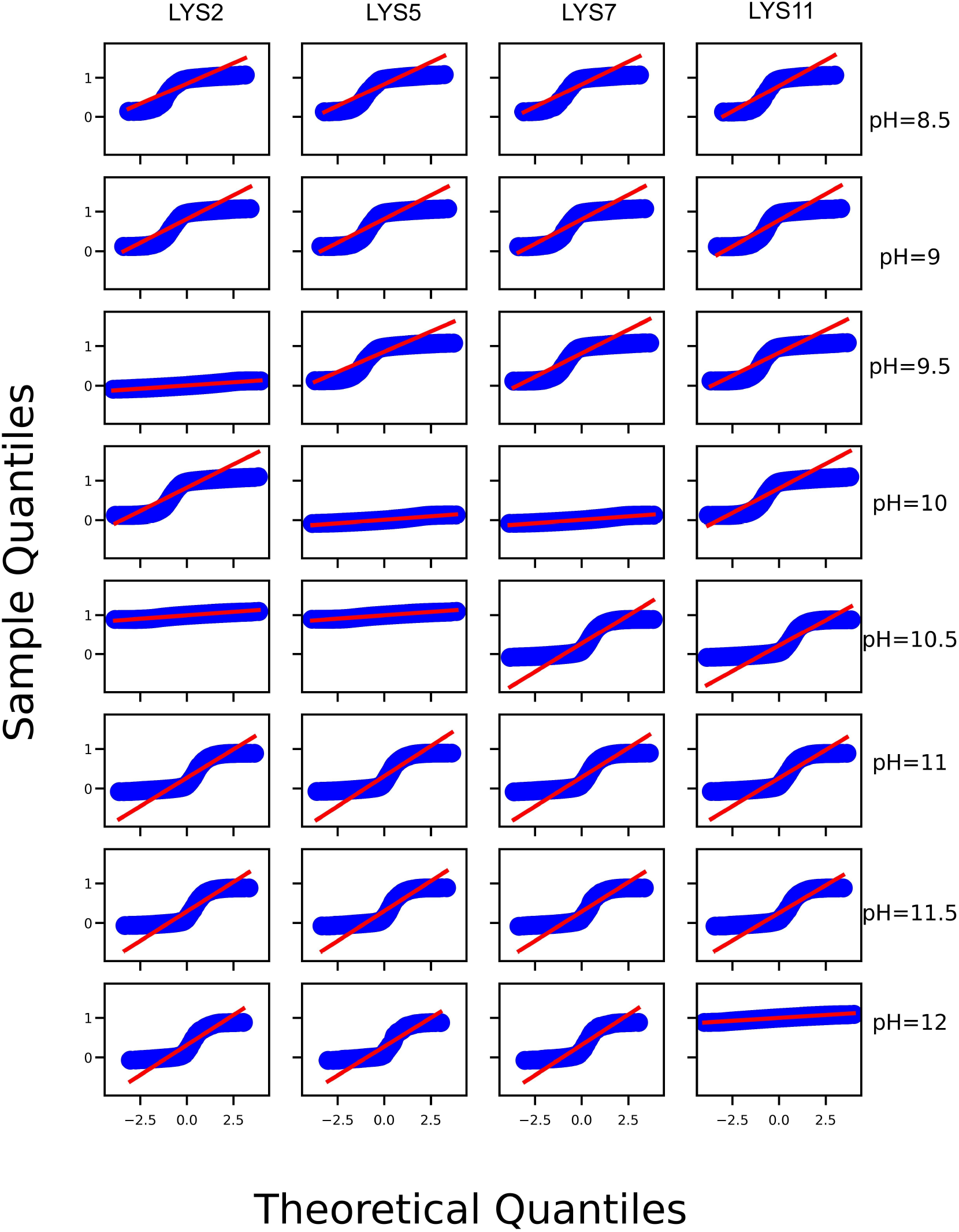
QQ plot, first mode of bimodal distribution function

**Fig. S3.**
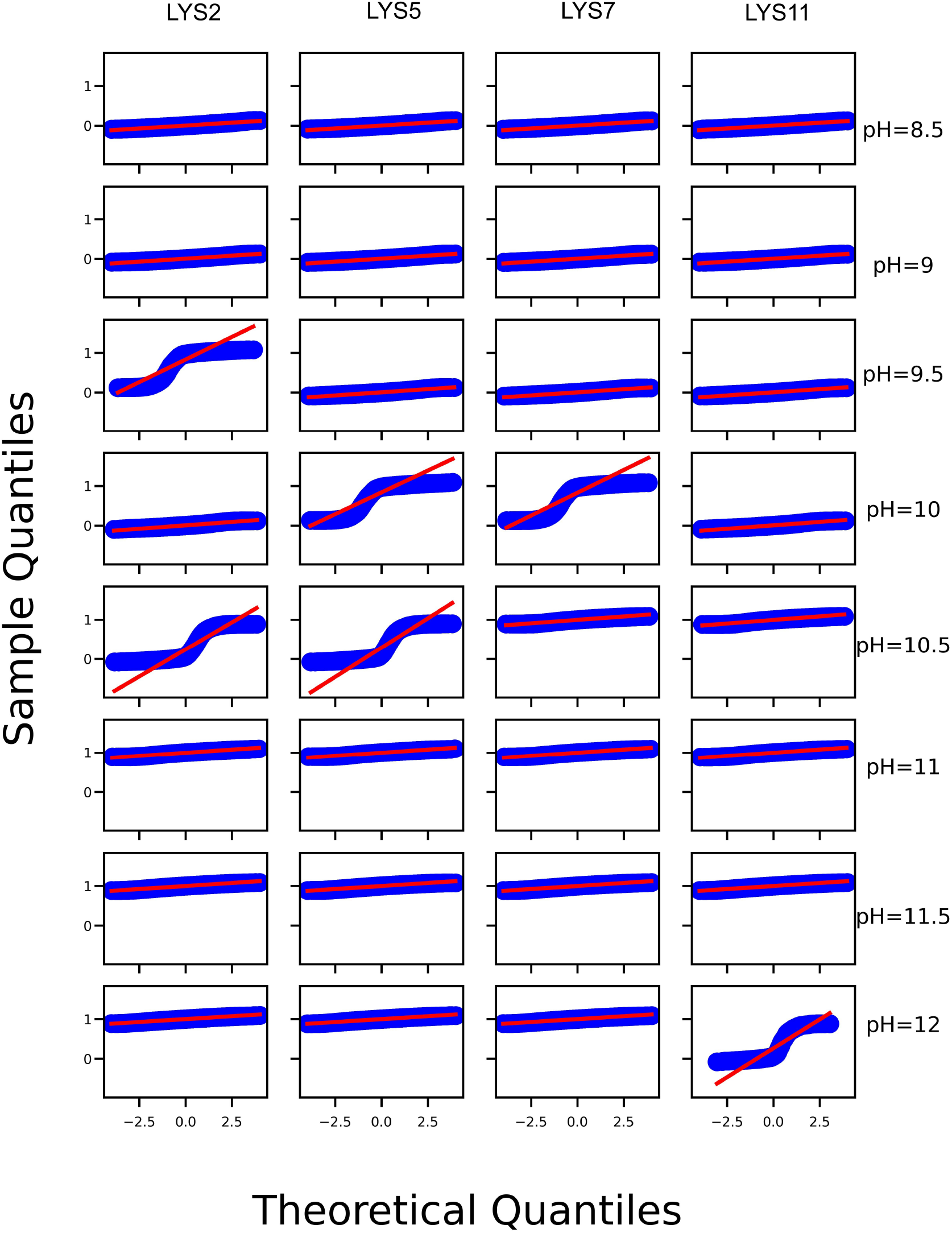
QQ plot, second mode of bimodal distribution function

**Fig. S4.**
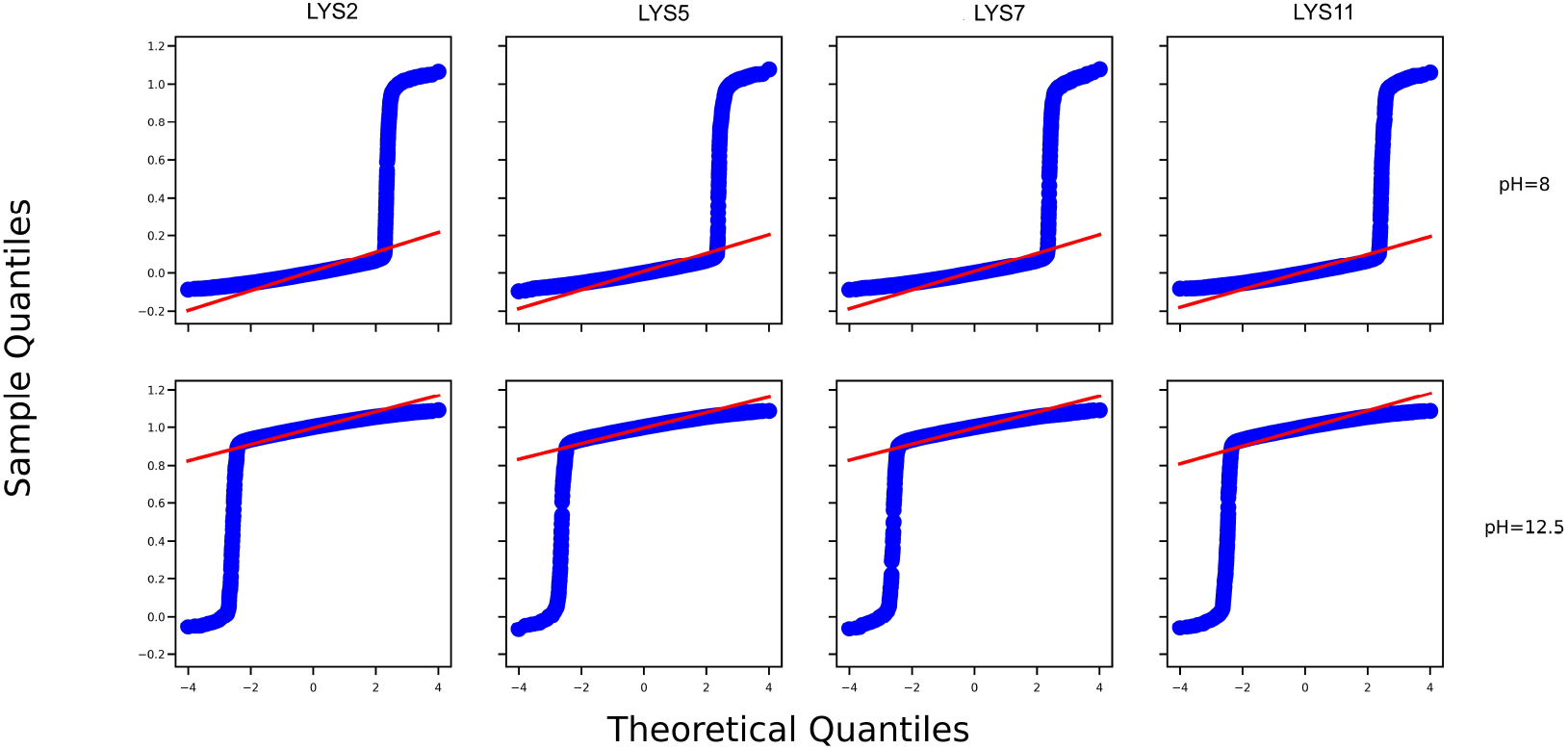
QQ plot, unimodal distribution function

**Table S2.**
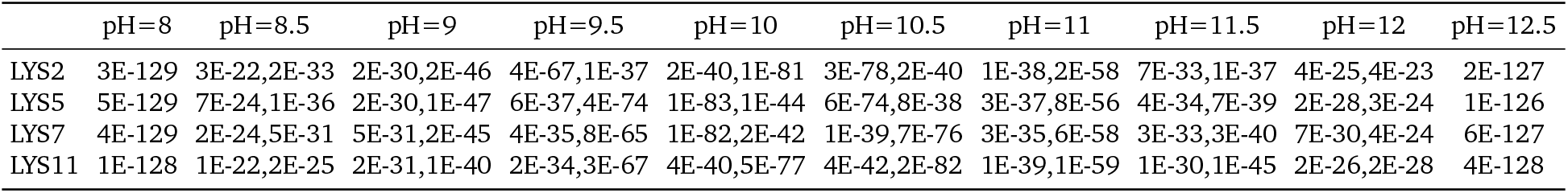
P-Values from Shapiro-Wilk test.

**Table S3.** P-Values from Levene Test.

|  | pH=8 | pH=8.5 | pH=9 | pH=9.5 | pH=10 | pH=10.5 | pH=11 | pH=11.5 | pH=12 | pH=12.5 |
| --- | --- | --- | --- | --- | --- | --- | --- | --- | --- | --- |
| LYS2 to LYS11 | 1E-2 | 4E-20 | 2E-22 | 1E-1 | 1E-40 | 1E-28 | 6E-3 | 1E-37 | 3E-2 | 3E-2 |
| LYS5 to LYS11 | 2E-1 | 5E-50 | 1E-26 | 8E-71 | 1E-90 | 5E-143 | 3E-12 | 9E-30 | 1E-1 | 6E-4 |
| LYS7 to LYS11 | 3E-1 | 2E-13 | 1E-16 | 3E-4 | 2E-53 | 1E-75 | 1E-3 | 3E-20 | 2E-1 | 5E-3 |

**Fig. S5.**
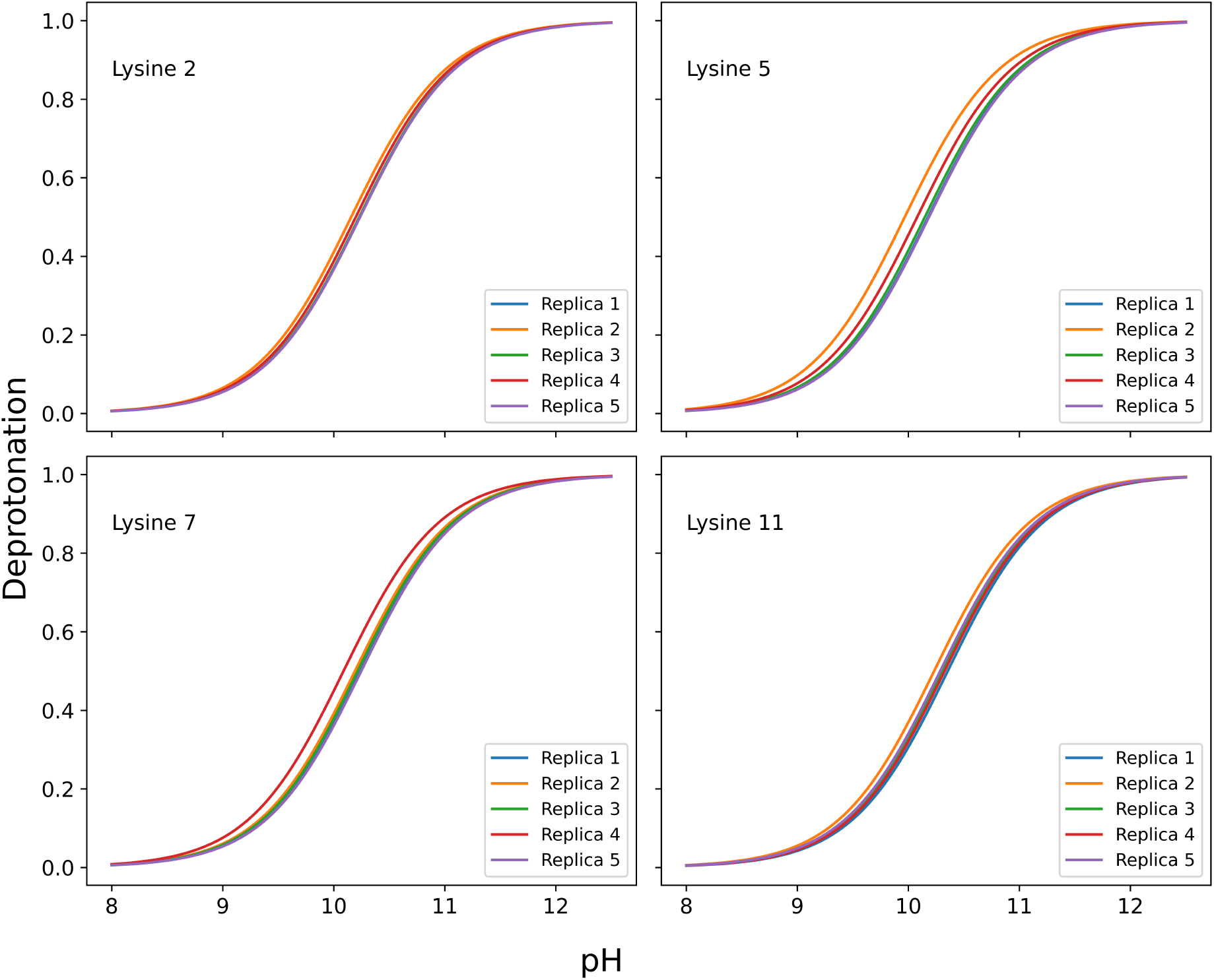
Calculated *pK*_*a*_ values of each lysine amino acid for each replica by fitting the deprotonated fraction into the Henderson-Hasselbach equation described in the methods section.

**Fig. S6.**
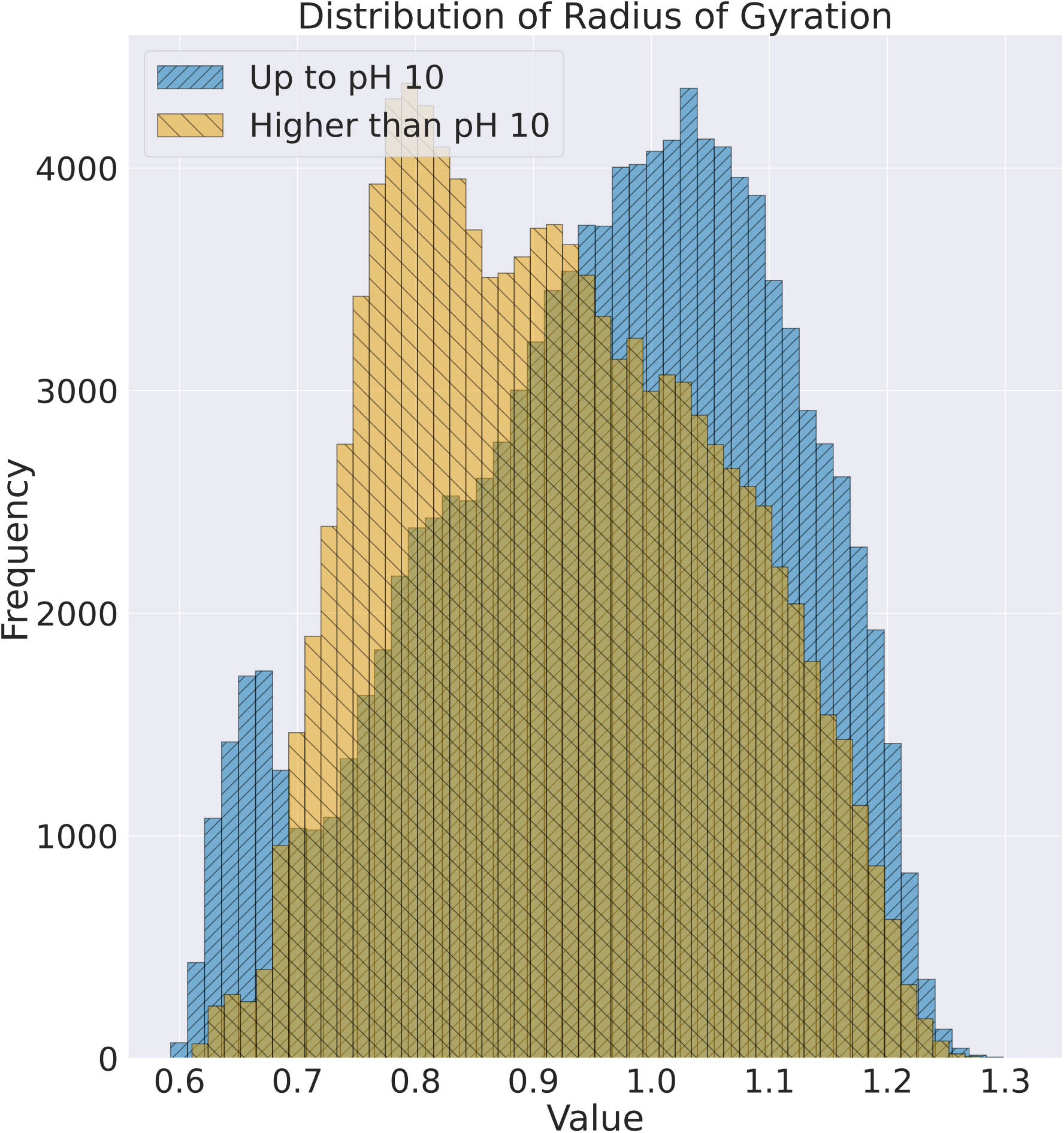
Distributions of radius of gyration for systems with pH value up to 10 and higher than 10.

**Fig. S7.**
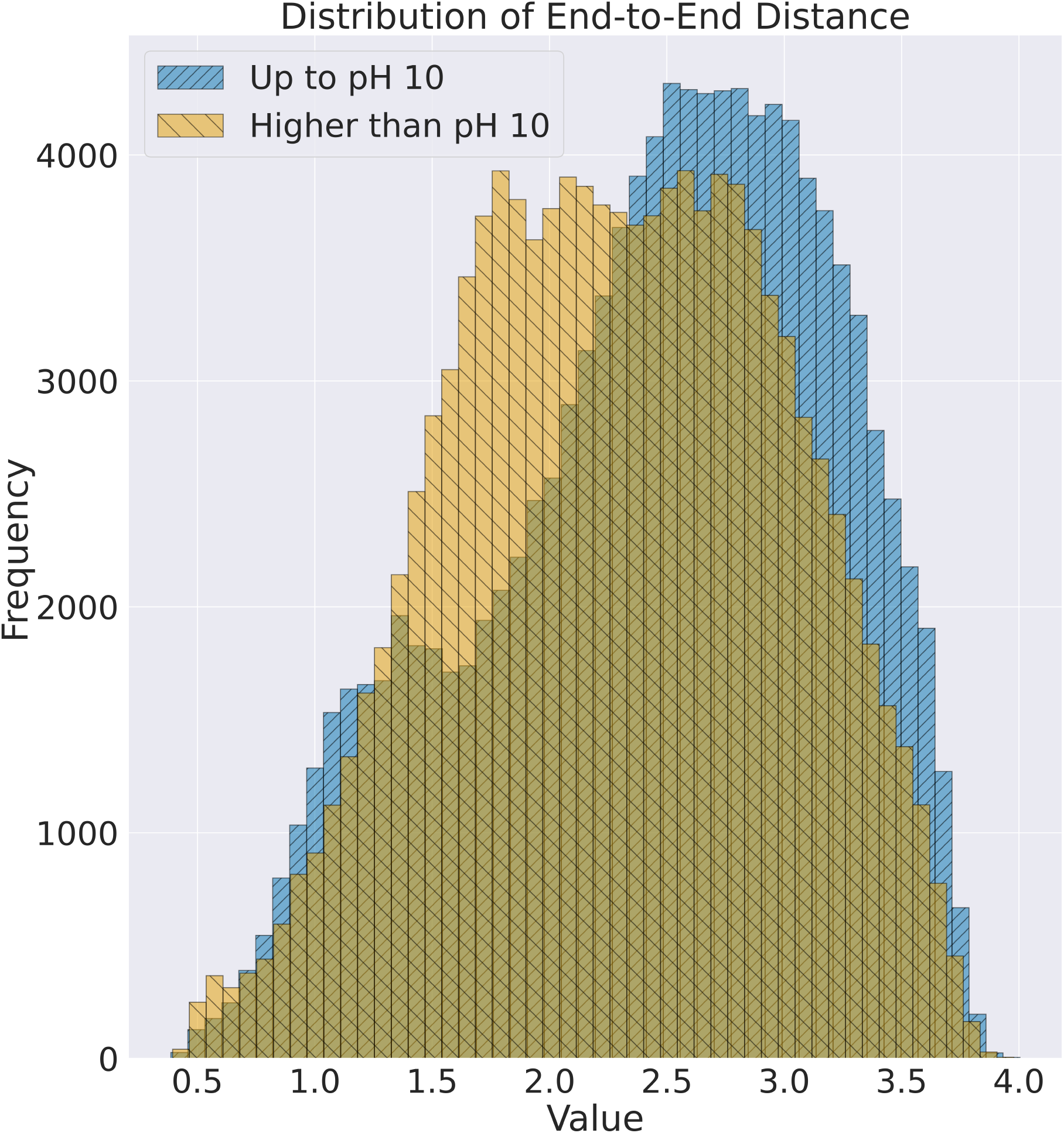
Distributions of End-to-End distances for systems with pH value up to 10 and higher than 10.

**Fig. S8.**
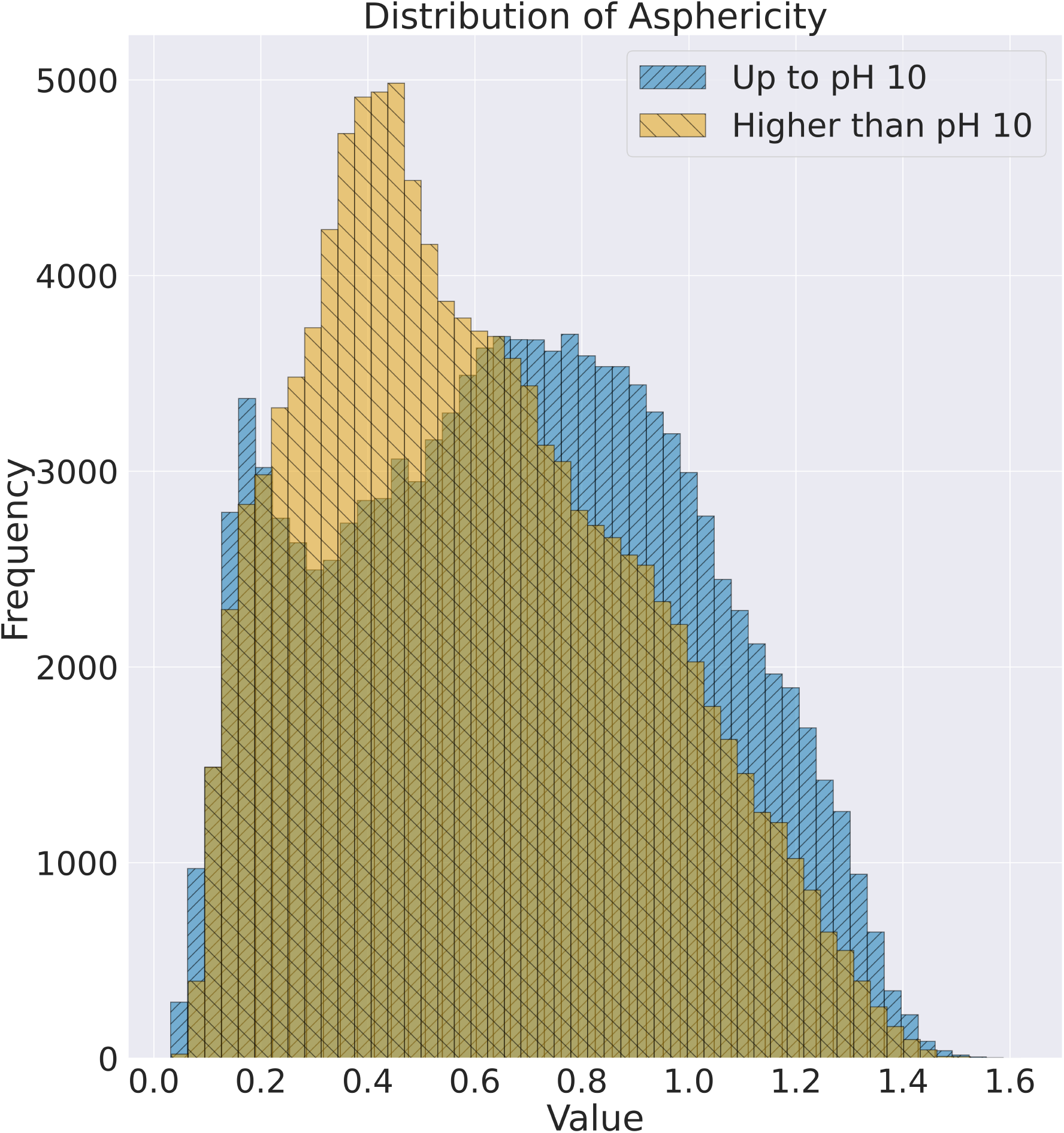
Distributions of asphericity for systems with pH value up to 10 and higher than 10.

**Fig. S9.**
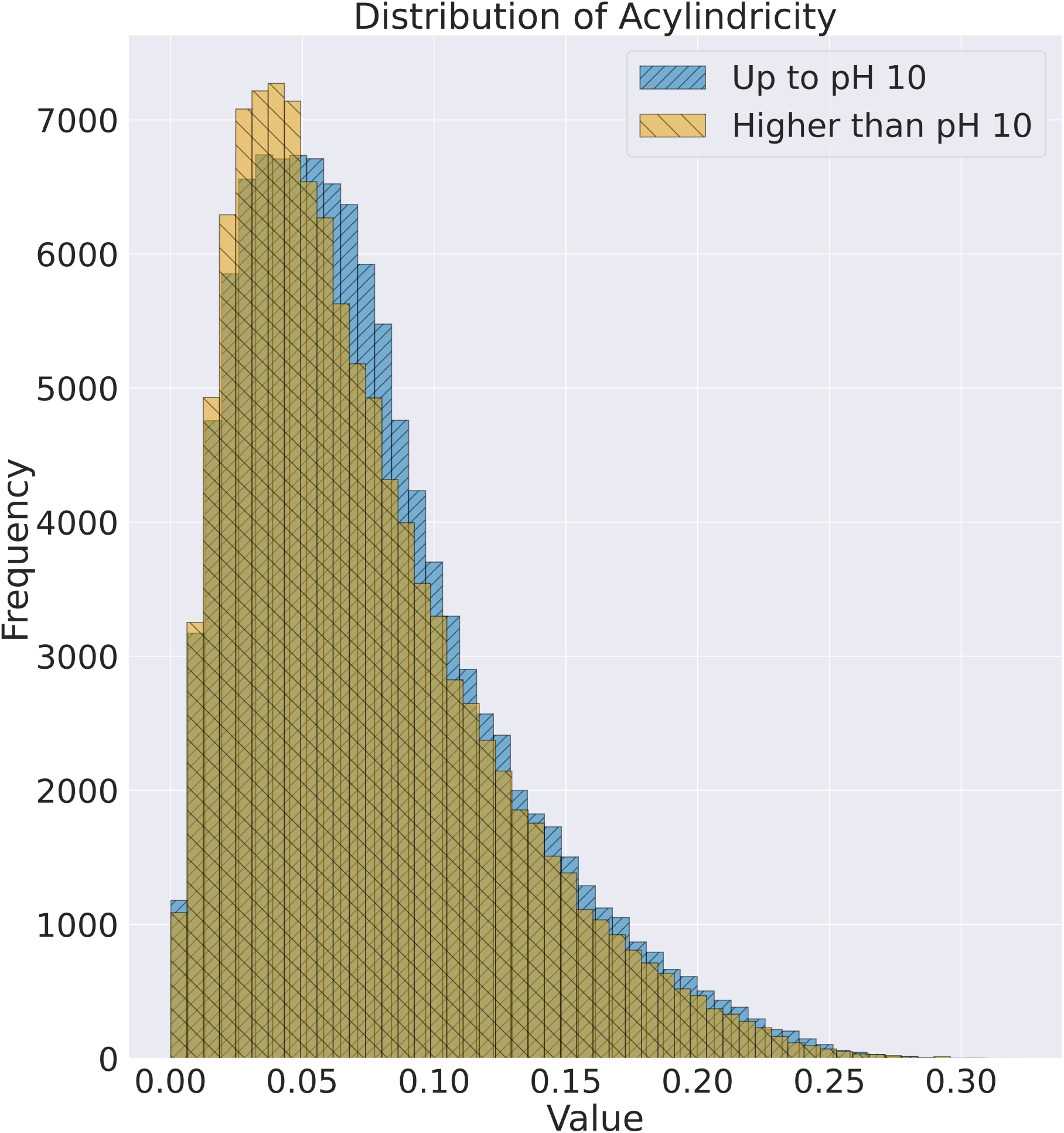
Distributions of acylindricity for systems with pH value up to 10 and higher than 10.

**Fig. S10.**
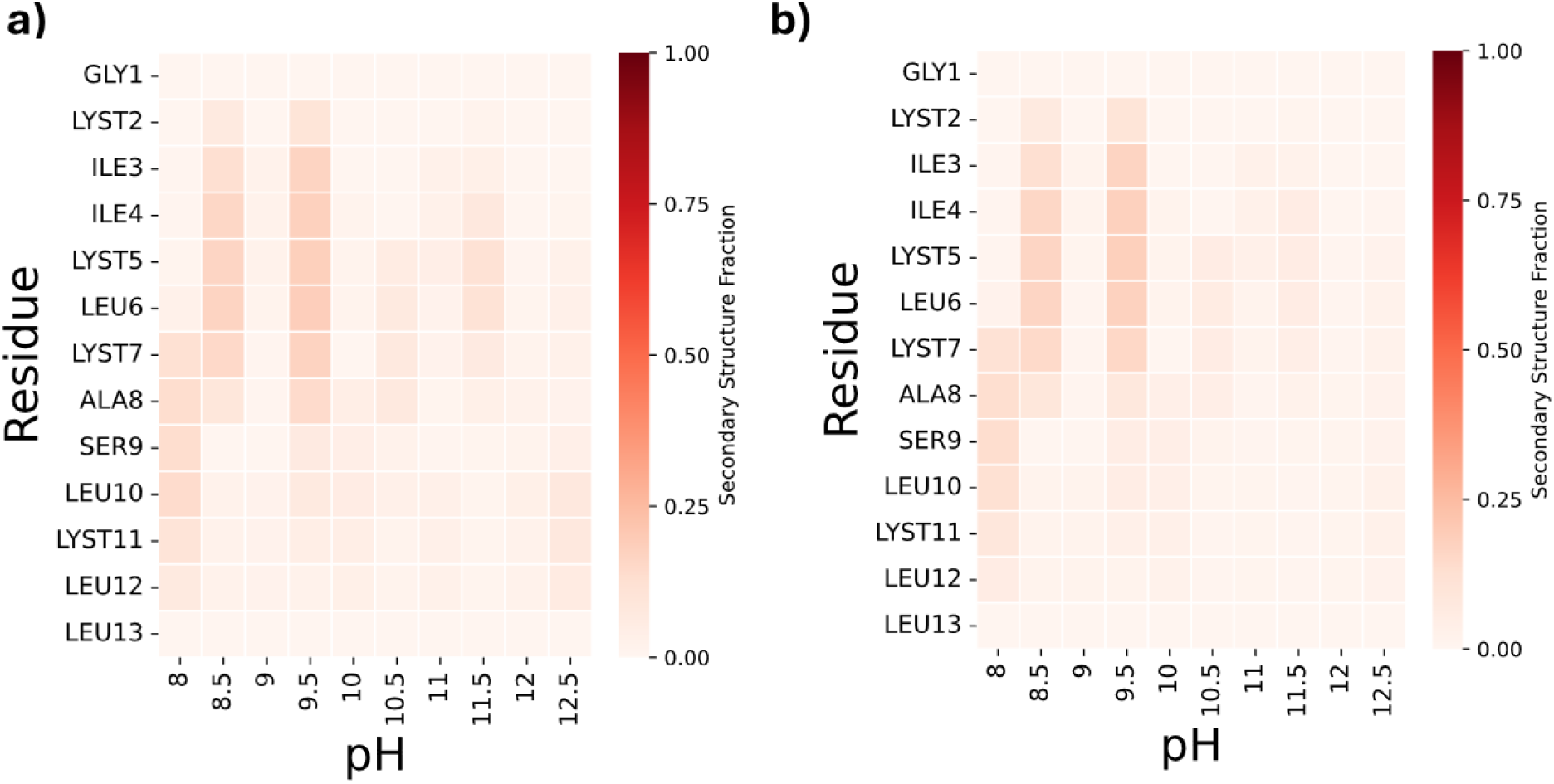
(a) Average helical structure per residue over 10 pH levels. (b) Error value per residue over 10 pH levels.

**Fig. S11.**
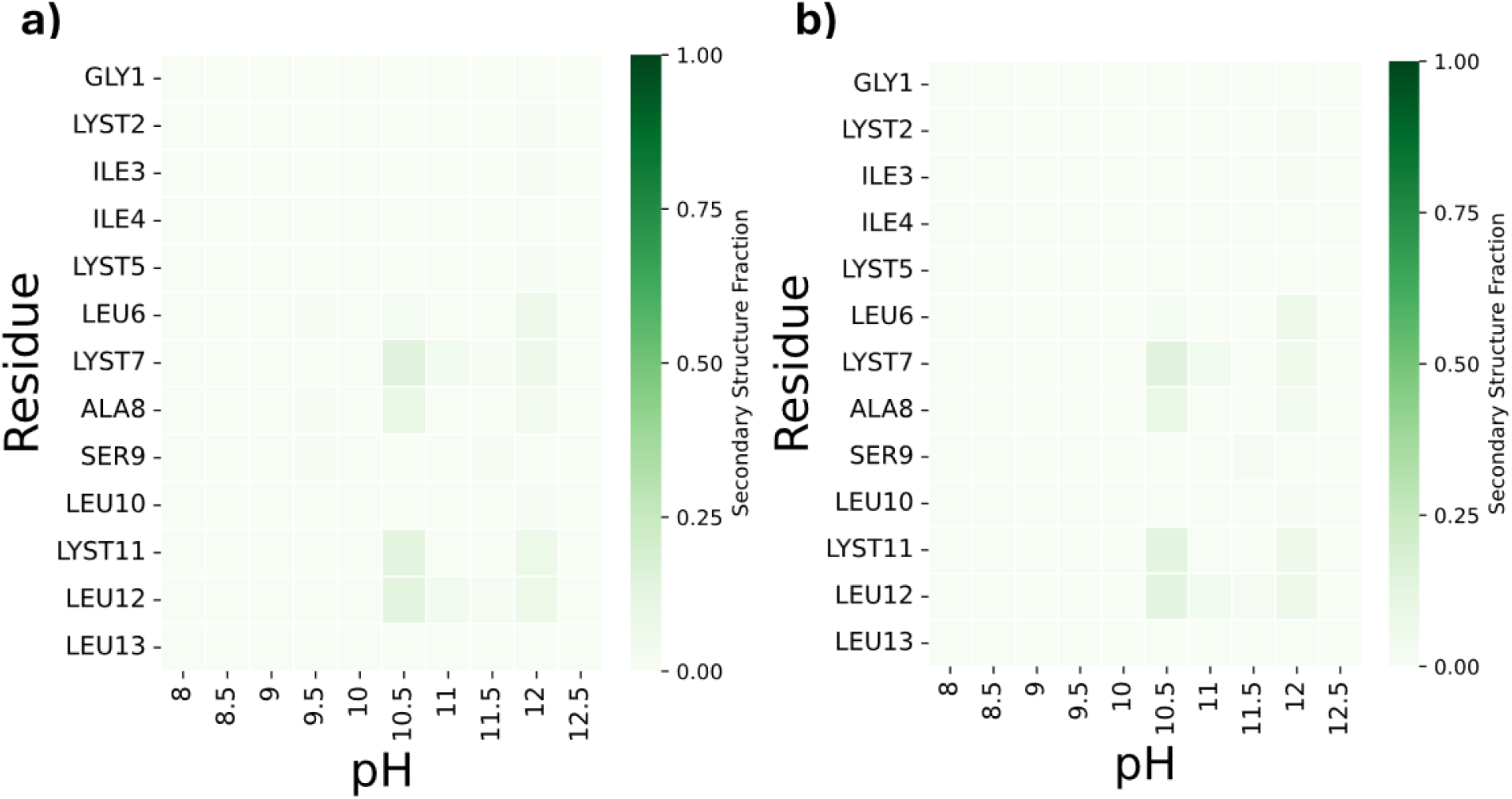
(a) Average strand structure per residue over 10 pH levels. (b) Error value per residue over 10 pH levels.

**Fig. S12.**
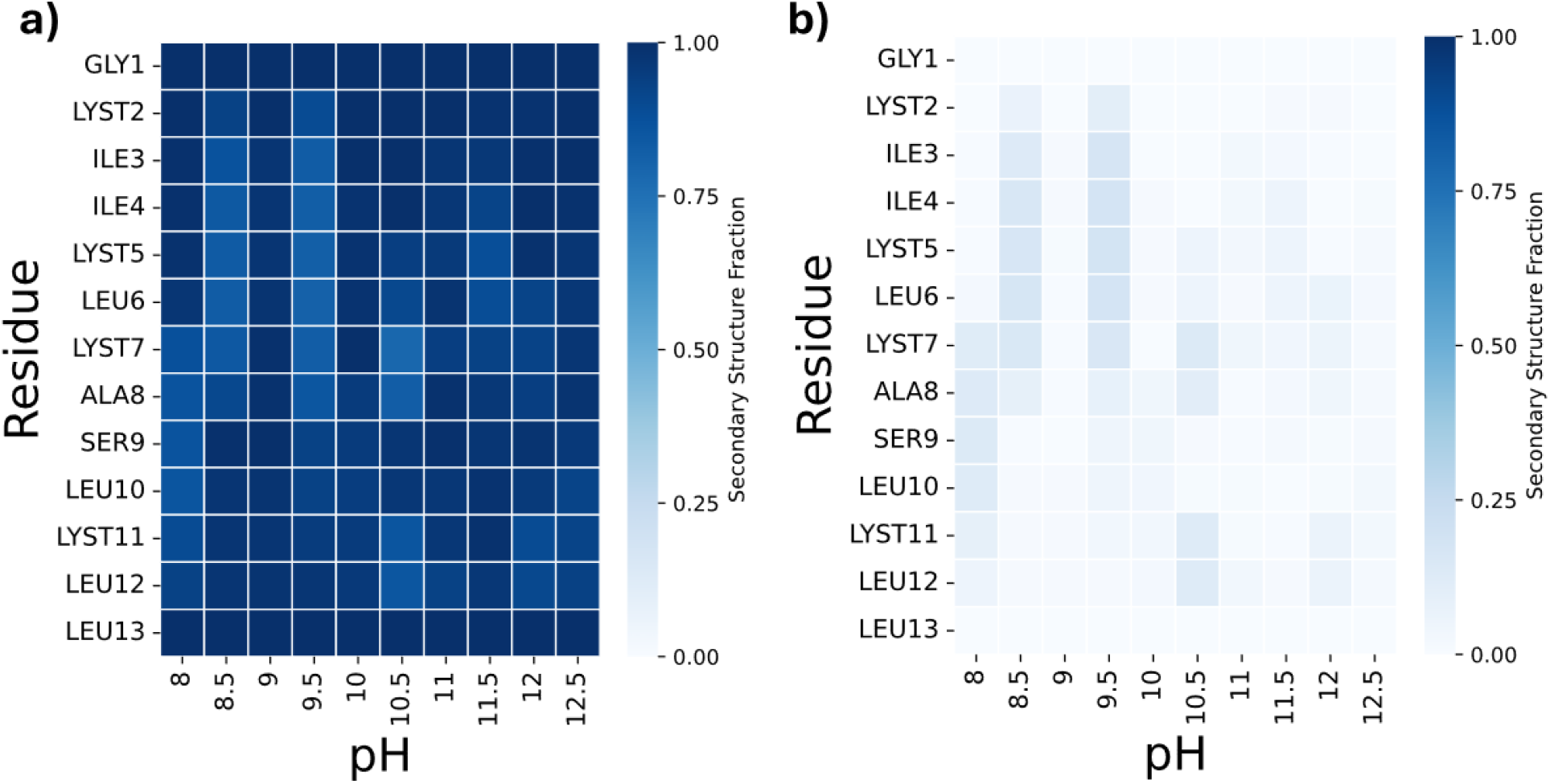
(a) Average coil structure per residue over 10 pH levels. (b) Error value per residue over 10 pH levels.

**Fig. S13.**
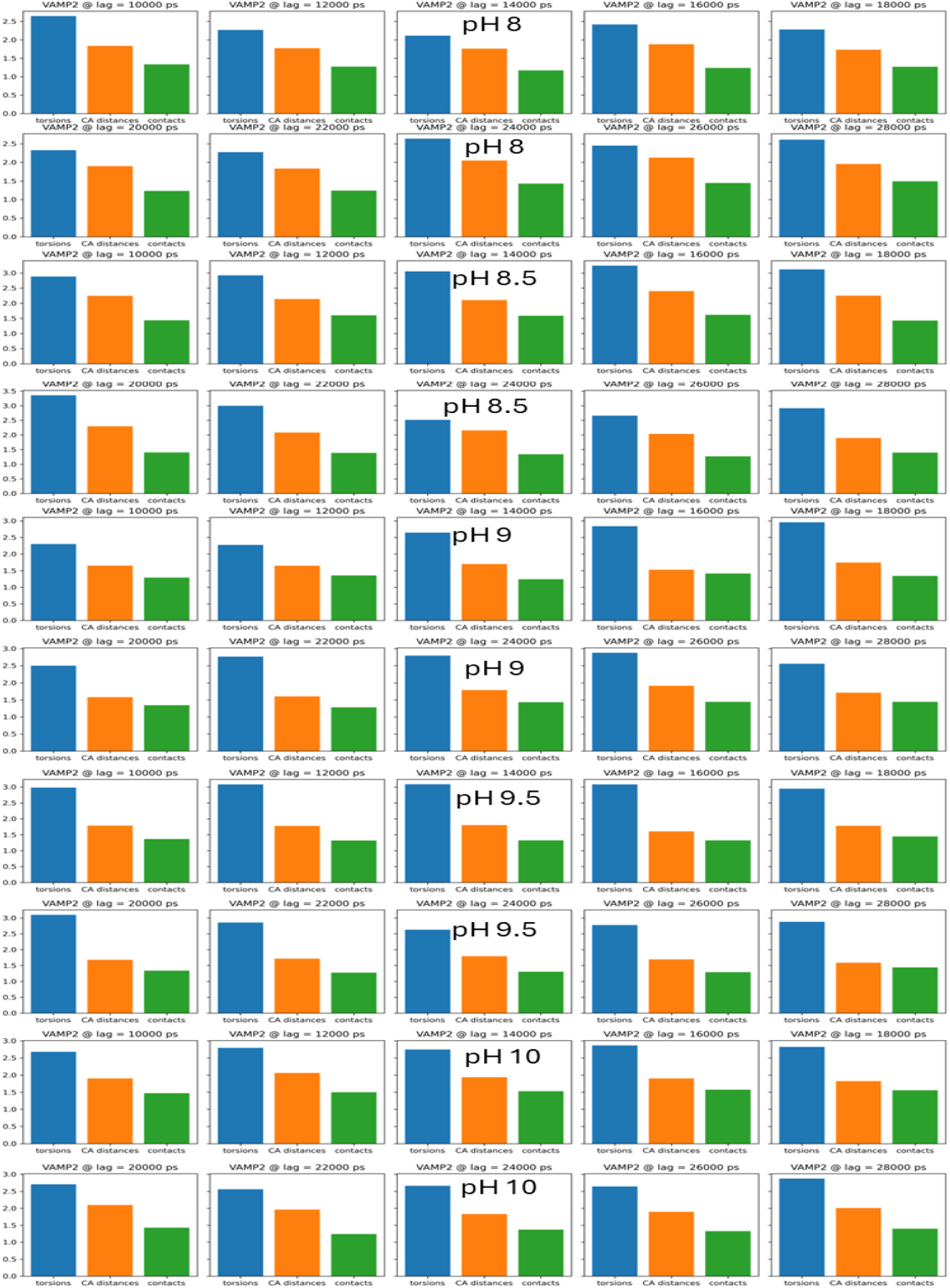
VAMP-2 score of individual systems from pH 8 to pH 10, calculated for 3 potential descriptors including the total dihedral angles (proper and improper), Alpha carbon (*C*_*α*_ ) distances, and non-hydrogen (heavy) atoms contacts at different lag times.

**Fig. S14.**
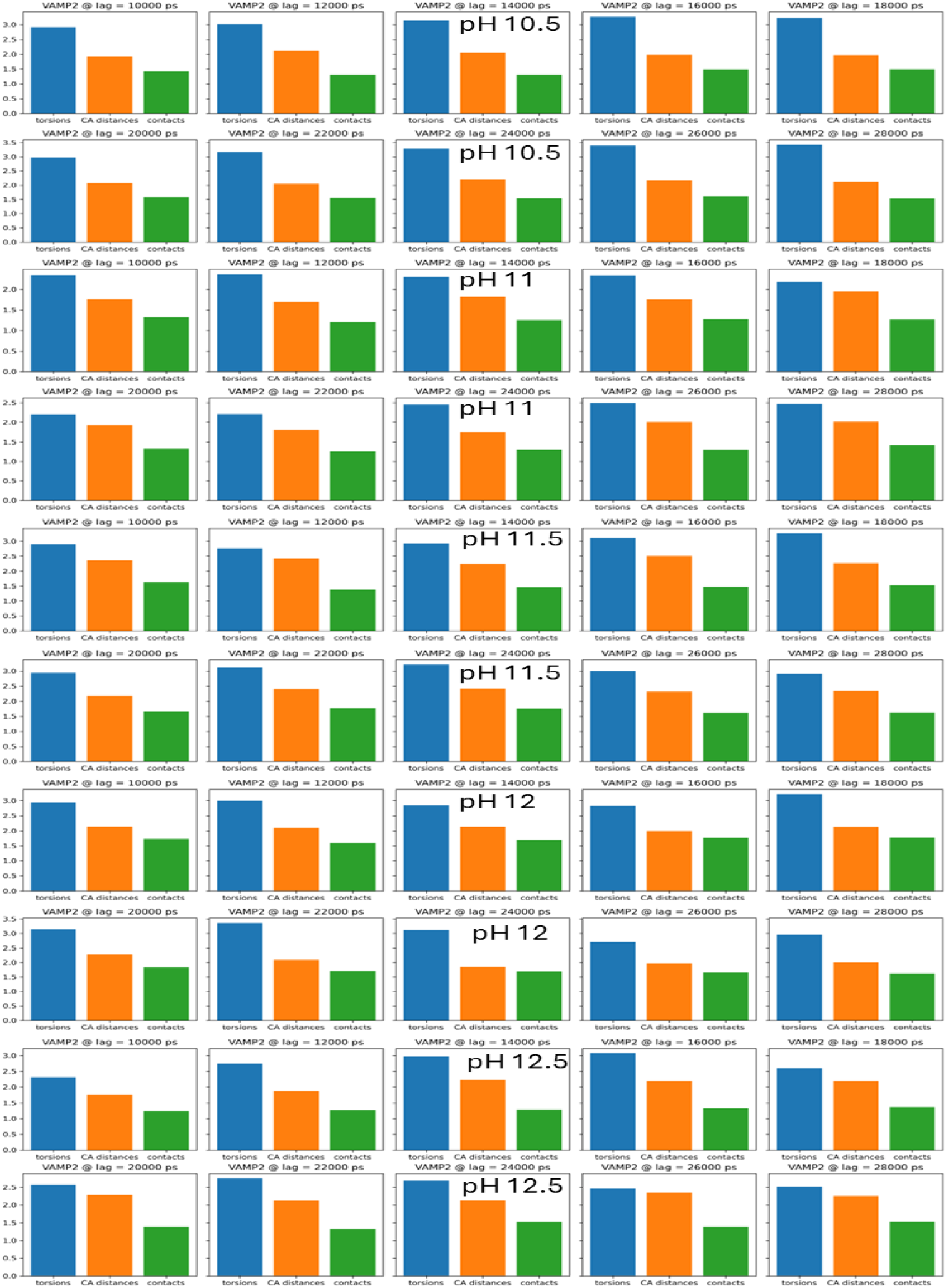
VAMP-2 score of individual systems from pH 10.5 to pH 12.5 calculated for 3 potential descriptors including the total dihedral angles (proper and improper), Alpha carbon (*C*_*α*_ ) distances, and non-hydrogen (heavy) atoms contacts at different lag times.

**Fig. S15.**
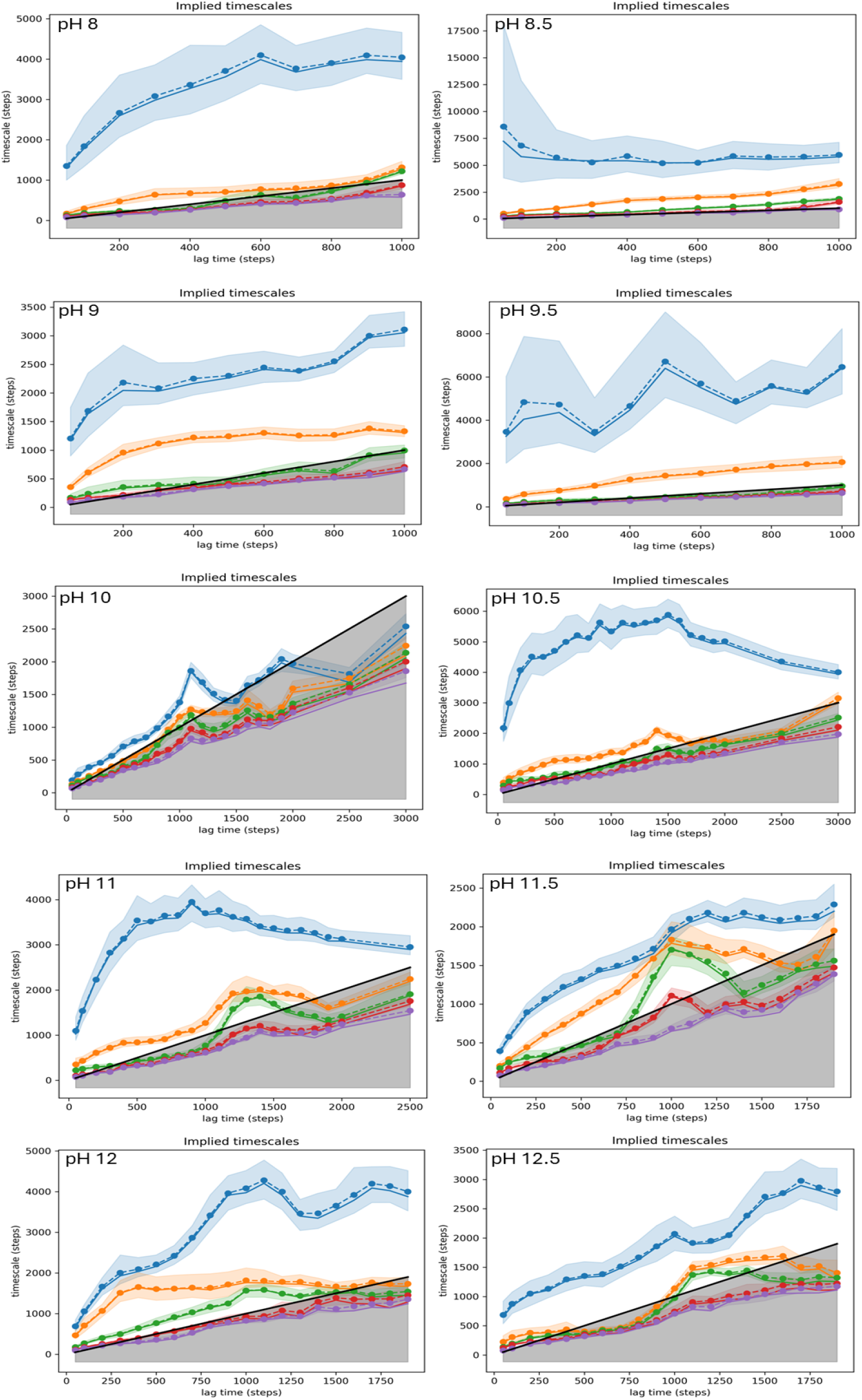
Separation of the implied timescales at various lag times at each pH levels for individual systems.

**Fig. S16.**
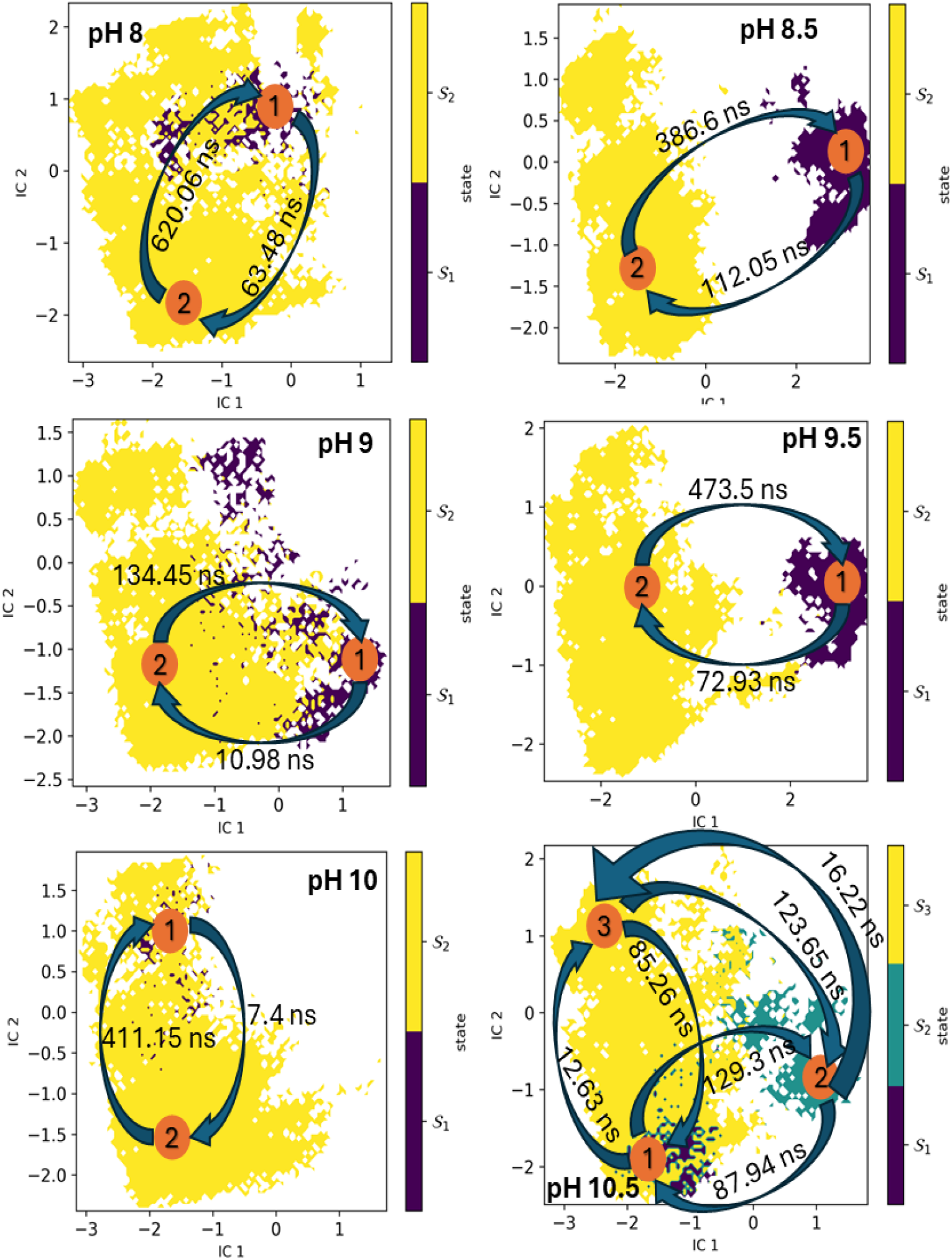
Free Energy Surface (FES) of the system under pH level of 8 to 10.5 computed using PCA as dimensionality reduction method, colored by metastable states, with mean first passage times (MFPTs) indicated for transitions between states.

**Fig. S17.**
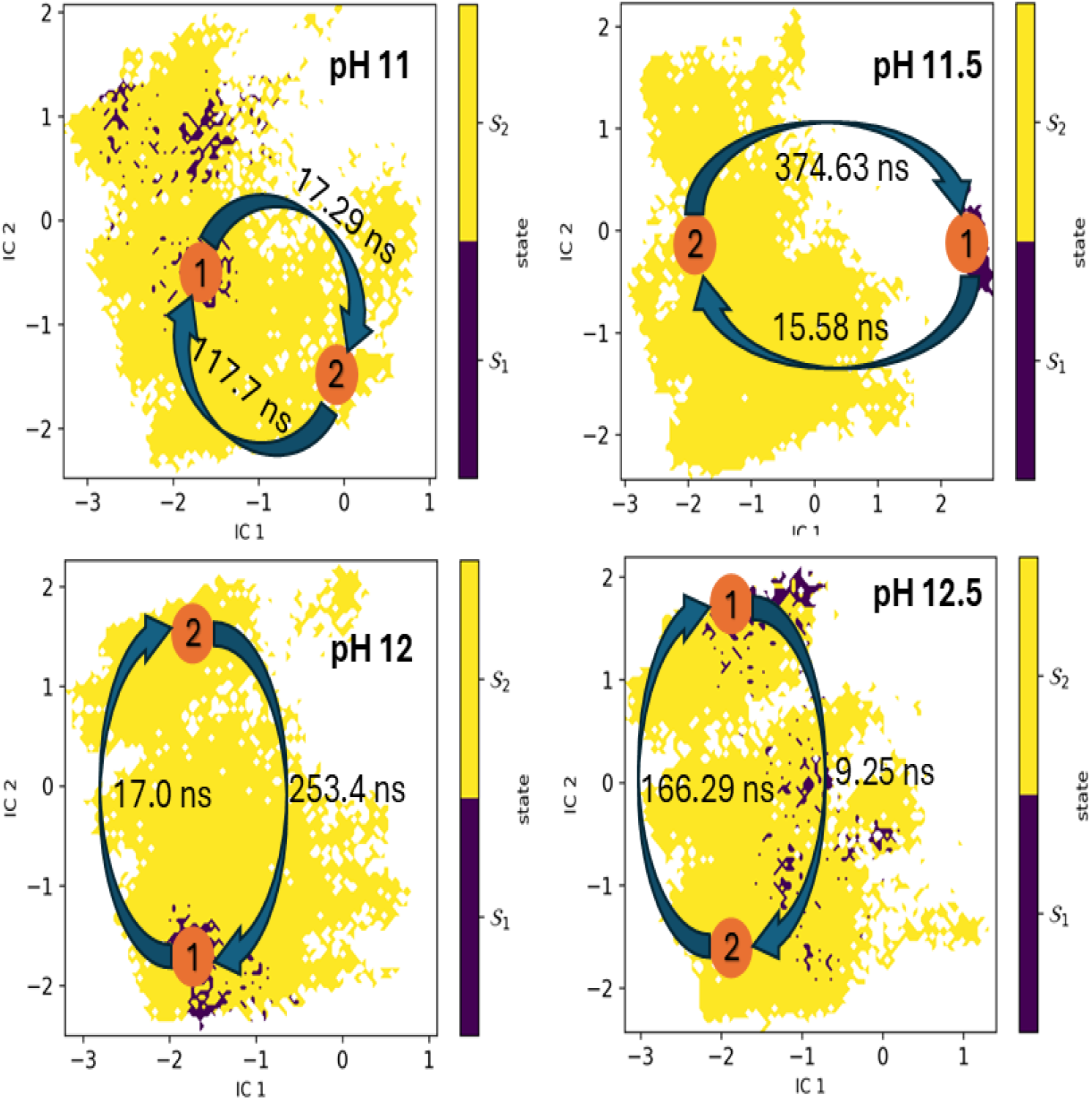
Free Energy Surface (FES) of the system under pH level of 11 to 12.5 computed using PCA as dimensionality reduction method, colored by metastable states, with mean first passage times (MFPTs) indicated for transitions between states.

**Fig. S18.**
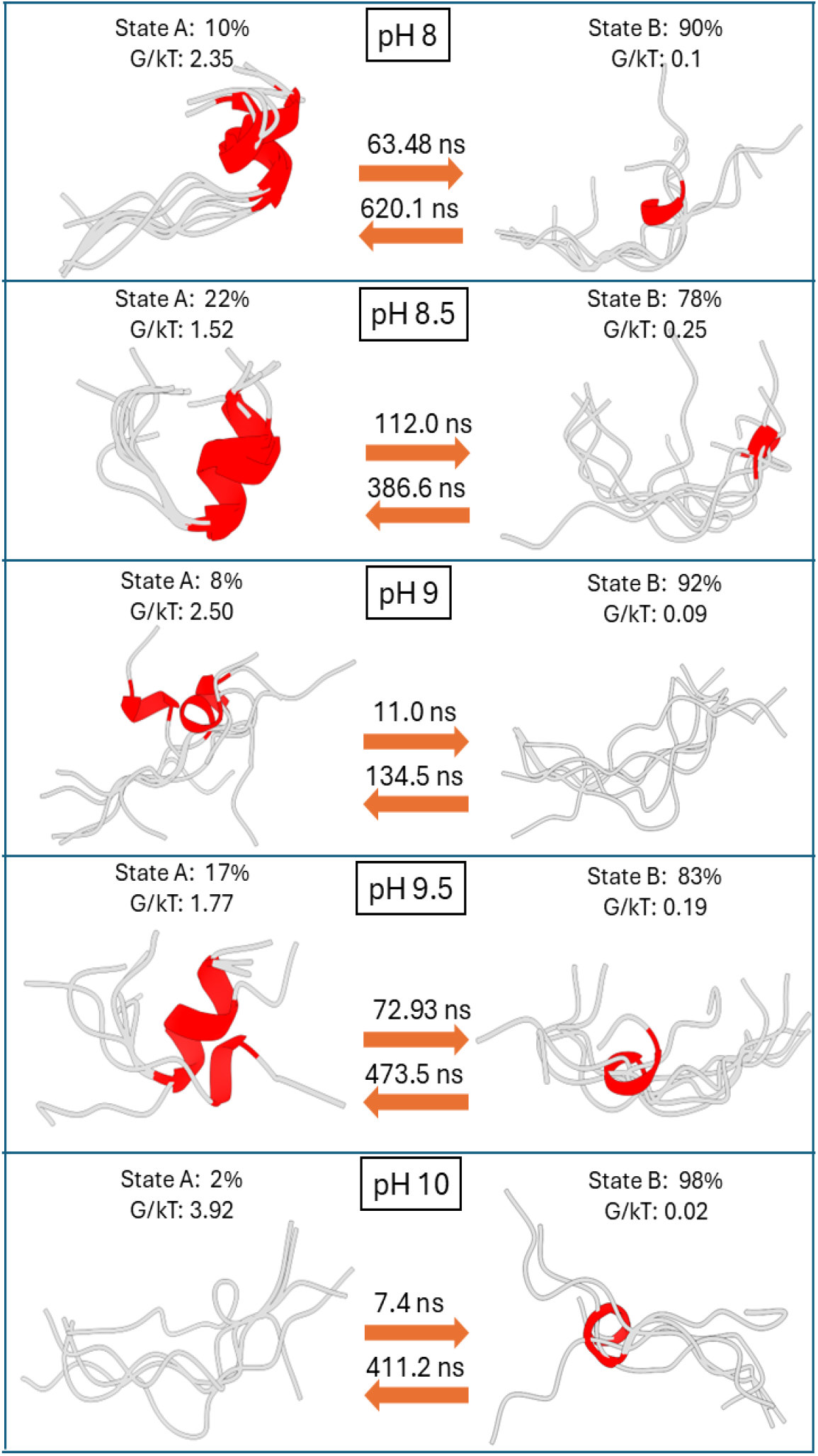
Metastable state population percentage of GL13K calculated by PCA dimensional reduction method from pH 8 to pH 10 is plotted with the representative structure of each state. These representative structures are based on the probability distributions of microstates identified by PCCA+. The mean first passage time (MFPT) between states is calculated in nanoseconds. The relative free energy for each state is shown under each stable state. The random coil and helical structures are colored in silver and red, respectively.

**Fig. S19.**
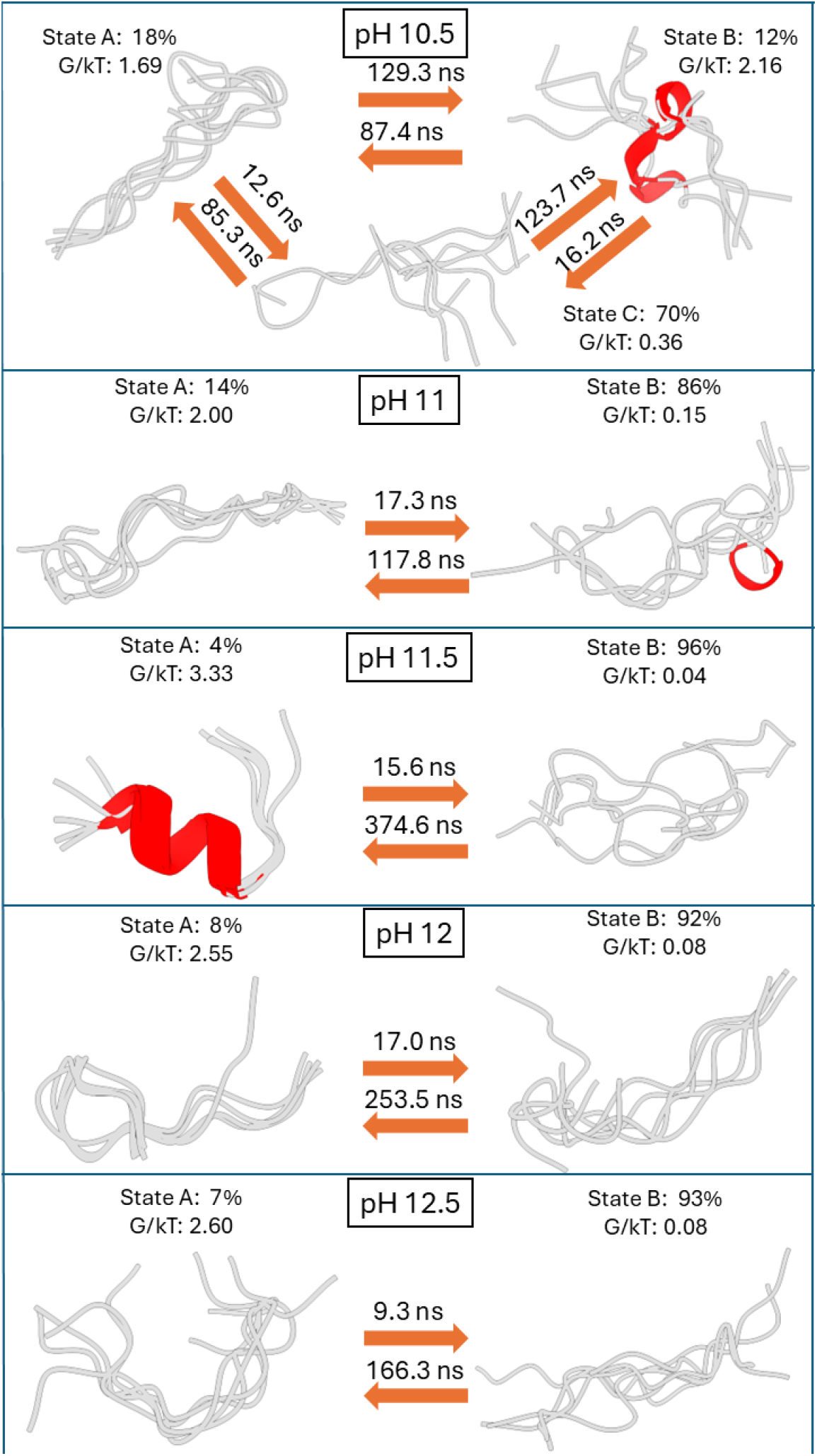
Metastable state population percentage of GL13K calculated by PCA dimensional reduction method from pH 10.5 to pH 12.5 is plotted with the representative structure of each state. These representative structures are based on the probability distributions of microstates identified by PCCA+. The mean first passage time (MFPT) between states is calculated in nanoseconds. The relative free energy for each state is shown under each stable state. The random coil and helical structures are colored in silver and red, respectively.

